# N-cadherin stabilises neural identity by dampening anti-neural signals

**DOI:** 10.1101/704817

**Authors:** K Punovuori, RP Migueles, M Malaguti, G Blin, KG Macleod, NO Carragher, T Pieters, F van Roy, MP Stemmler, S Lowell

## Abstract

A switch from E-to N-cadherin regulates the transition from pluripotency to neural identity but the mechanism by which cadherins regulate differentiation was previously unknown. Here we show that the acquisition of N-cadherin stabilises neural identity by dampening anti-neural signals. We use quantitative image-analysis to identify an effect of N-cadherin to promote neural differentiation independently of effects on cell cohesiveness. We reveal that cadherin switching diminishes the level of nuclear β-catenin, and that N-cadherin also dampens FGF activity and consequently stabilises neural fate. Finally, we compare the timing of cadherin switching and differentiation *in vivo* and *in vitro*, and find that this process becomes dysregulated during *in vitro* differentiation. We propose that N-cadherin helps to propagate a stable neural identity throughout the emerging neuroepithelium, and that dysregulation of this process contributes to asynchronous differentiation in culture.

## Introduction

There is an increasing appreciation that changes in adhesion and morphology help to regulate cell fate changes (1). The homotypic adhesion molecule E-cadherin is expressed on the surface of pluripotent cells and is downregulated and replaced with N-cadherin during early neural development (2,3). We previously reported that loss of E-cadherin is not simply a consequence of differentiation, but rather that it actively promotes the neural differentiation process (4). However the role of N-cadherin in this process and the mechanisms by which cadherins regulate neural differentiation are not known.

It has previously been reported that premature cadherin switching has profound effects at gastrulation, including an expansion of the extraembryonic compartment, a reduction in the size of the epiblast, and mis-patterning of the germ layers (5). These diverse phenotypes can be attributed at least in part to an overall reduction in BMP signalling within the epiblast and a reduction in pro-mesoderm signals at the primitive streak, which in turn may result from the gross morphological defects seen in these embryos (5). However it is not clear which aspects of this complex phenotype are an indirect consequence of defects in extraembryonic tissues and which, if any, are cell-autonomous. Here we use cultured mouse pluripotent cells in order to focus on the mechanism by which cadherin switching influences neural differentiation of pluripotent cells in the absence of extraembryonic tissues.

Mouse pluripotent cells can be cultured the presence of inhibitors of MEK and Gsk3β plus LIF (2i+LIF) in order to maintain them in a naïve embryonic stem cell (ESC) state equivalent to the preimplantation epiblast (6,7), or they can be cultured in the presence of FGF and Activin in order to maintain a differentiation-primed epiblast stem cell (EpiSC) state equivalent to the postimplantation epiblast (8–10). LIF and foetal calf serum (FCS) support a heterogeneous mixture of pluripotent cells moving in and out of the naïve state. We previously reported that cells downregulate E-cadherin during neural differentiation of ES cells, and that loss of E-cadherin leads to faster, more synchronous neural differentiation in vitro (4), in keeping with other reports that E-cadherin acts as a ‘brake’ to slow down differentiation of pluripotent cells (11–16). E-cadherin-null ESCs display a loss of cell-cell adhesion (17,18), raising the possibility that their neural differentiation phenotype may be a secondary consequence of their adhesion defect. Alternatively, cadherins could influence differentiation by modulating signalling independently of adhesion (2,14,19,20).

Neural specification depends on inhibition of BMP and Nodal signalling (21,22). The ability of BMP to block neural fate is at least in part due to maintenance of E-cadherin expression, but it is not know which signalling pathways act downstream of cadherins to modulate differentiation. Dampening of either FGF (23–26) or Wnt (27,28) has the effect of stabilising neural identity. N-cadherin has been reported to modulate FGF activity (29–32) and E-cadherin has been reported to modulate Wnt activity in other contexts (33), and so it seems plausible that cadherin switching may modulate neural differentiation via dampening of one or both of these anti-neural signalling pathways. Alternatively, it is possible that cadherins modulate other signalling pathways (34).

Here, we set out to ask how the switch from E-cadherin to N-cadherin influences differentiation. We present evidence that N-cadherin promotes neural differentiation by dampening FGF activity. We also discover that cadherin switching occurs later and more synchronously during anterior neural differentiation *in vivo* compared with neural differentiation in culture. We suggest that cadherins could mediate a ‘community effect’ by helping to propagate differentiation decisions to neighbouring cells, and that this may help to ensure synchronous neural commitment in the embryo. This effect partly breaks down in culture, helping to explain why differentiation in culture is relatively asynchronous even in the face of a uniform extrinsic environment.

## Results

### Cadherin switching is initiated prior to the onset of neural differentiation *in vitro*

We previously reported that E-cadherin inhibits neural differentiation, but the mechanism of action was not known (4). Upregulation of N-cadherin accompanies the loss of E-cadherin as pluripotent cells adopt a neural fate (3,35), raising the possibility that N-cadherin might contribute to the regulation of the differentiation process. We first asked when N-cadherin becomes detectable during neural differentiation.

We confirmed that mouse embryonic stem cells cultured in 2i-Lif or Lif-serum express high levels of E-cadherin, whereas N-cadherin mRNA and protein were undetectable in either of these culture conditions (Fig. 1A-B). In EpiSC culture, E-cadherin expression was heterogeneous while N-cadherin became detectable in a subpopulation of cells (Fig. 1A-C). Cultures of EpiSCs contain spontaneously differentiating cells, and so we focused only on undifferentiated (Oct4^+^) cells (Fig. 1D, Fig. S1A). Almost all (99.6%) Oct4^+^ cells expressed E-cadherin and, of these, 13.0% also expressed N-cadherin (Fig. 1E). Very few (<1%) Oct4^+^ cells expressed N-cadherin alone. These results show that N-cadherin becomes expressed in a subpopulation of E-cadherin^+^ cells prior to loss of Oct4 expression.

**Figure 1:**
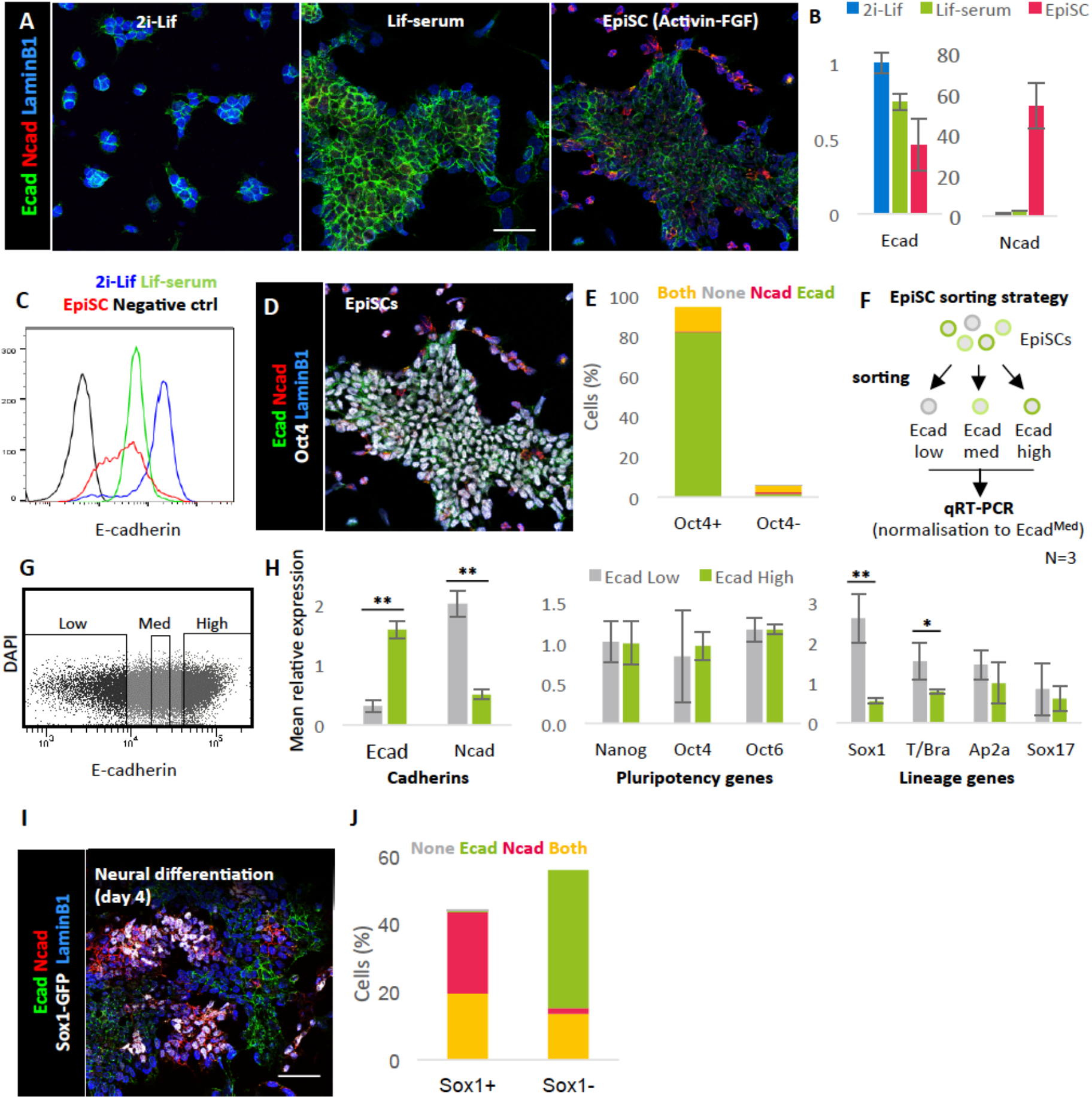
Cadherin switching precedes the loss of pluripotency marker expression and coincides with neural priming *in vitro*. **A.** Cells cultured in three pluripotent conditions stained for E-cadherin, N-cadherin, and the nuclear envelope marker LaminB1. **B.** qRT-PCR analysis of E-cadherin expression in cells cultured in three pluripotent conditions, N=3. **C.** Flow cytometric analysis of E- and N-cadherin expression in cells cultured in various pluripotent conditions; curves show representative sample of three biological replicates. **D.** Example ICC image of EpiSCs stained Ecad, Ncad, Oct4 and LaminB1 **E.** Quantification of protein co-expression in EpiSCs. N=2596 cells from three biological replicates. **F.** Sorting and analysis strategy for EpiSCs heterogeneously expressing E-cadherin. **G.** Example FACS gating of EpiSCs into three populations based on their level of E-cadherin expression; each population makes up ~20% of live cells. **H.** qPCR analysis of sorted EpiSC populations. Ecad^Low^ and Ecad^High^ populations were normalised to the Ecad^Med^ population. N=3 independent sorts. **I.** Example ICC image of Sox1-GFP (46C) cells undergoing neural differentiation stained for Ecad, Ncad, Sox1 and Lamin1. **J.** Quantification of protein co-expression in Sox1-GFP cells undergoing neural differentiation. N=2275 cells from three biological replicates. Error bars=SD, *p≤0.05, **p≤0.01, paired T-test. All images shown to same scale, scale bars=50um.

FACS analysis confirmed that the vast majority of EpiSCs express E-cadherin, but revealed considerable cell-to-cell variability in the levels of this adhesion molecule on the cell surface (Fig. 1C, red curve). This contrasted with naïve pluripotent cells, which displayed uniformly high levels of E-cadherin throughout the population (Fig 1C, blue curve). EpiSCs were sorted into three subpopulations (Ecad^High^, Ecad^Med^, and Ecad^Low^)(Fig. 1F-G). Ecad^High^ and Ecad^Low^ populations were then analysed by qRT-PCR, normalising to the Ecad^Med^ population (Fig. 1H). This analysis revealed a reciprocal expression pattern between E-cadherin and N-cadherin, consistent with an ongoing process of cadherin switching (Fig. 1H). The subpopulations expressed similar levels of the general pluripotency factors Oct4 and Nanog, and the primed pluripotency factor Oct6, indicating no difference in their overt differentiation status (Fig. 1H). It has been reported that differentiation-primed subpopulations of undifferentiated EpiSC express low levels of either the neural-priming factor Sox1 or the mesoderm-priming factor T/Brachyury (36). We found that the E-cadherin^Low^ population expressed significantly higher levels of Sox1 and T/Brachyury, characteristic of a differentiation-primed subpopulation, while markers of surface ectoderm (Ap2a) or endoderm (Sox17) did not differ significantly between the populations (Fig. 1H).

These results indicate that N-cadherin starts to become detectable prior to the loss of pluripotency transcription factors, and that a subpopulation of EpiSCs with lower E-cadherin and higher N-cadherin may be primed for neural and mesodermal differentiation.

We next examined cultures at an early stage of neural differentiation (d4), where around 50% of cells had started to adopt a neural identity, as revealed by immunostaining for Sox1-GFP (37), a reporter for the earliest marker of neuroepithelial identity (38) (Fig. 1I, Fig. S1B). In these cultures, N-cadherin was detectable in almost all Sox1-GFP^+^ neural cells, and of these around half also retained E-cadherin expression. Of cells that had not yet acquired a Sox1-GFP^+^ neural identity, almost all expressed E-cadherin, and of these around 20% also expressed N-cadherin (Fig. 1J).

Taken together, these results suggest that cadherin switching is initiated prior to the onset of neural differentiation *in vitro*.

### N-cadherin promotes neural differentiation

Loss of E-cadherin promotes neural differentiation of ESCs by an unknown mechanism (4). One possibility is that this pro-neural differentiation phenotype of cells lacking E-cadherin function may be a secondary consequence of their adhesion defect (Larue et al 1994., Larue et al., 1996). If this is the case, then restoring adhesion should restore normal neural differentiation capacity. N-cadherin can rescue the E-cadherin adhesion phenotype, at least in the context of preimplantation development (19,39). We therefore asked whether providing N-cadherin to E-cadherin null cells would rescue their neural differentiation phenotype.

To address this question, we made use of two E-cadherin knockout ES cell lines: Ecad^-/-^, where both cadherin alleles have been knocked out (40), and an N-cadherin knock-in line Ecad^Ncad/Ncad^, in which the coding sequence of N-cadherin is knocked into the E-cadherin locus, placing exogenous N-cadherin under the control of the endogenous E-cadherin regulatory elements and eliminating E-cadherin expression (5,39,41)(Fig. 2A). The N-cadherin knock-in (Ecad^Ncad/Ncad^) cells have been previously shown to enforce a cadherin switch while maintaining cell-cell adhesion in some developmental contexts (39). Since the two lines differ in their genetic background, they are analysed side-by-side with their relevant control cell lines, Ecad^Flox/Flox^ and Ecad^WT/WT^, respectively. We confirmed that N-cadherin was expressed above control levels in Ecad^Ncad/Ncad^ cells throughout the course of differentiation, but were not significantly elevated above background levels in Ecadh^-/-^ cells (Fig 2B).

**Figure 2:**
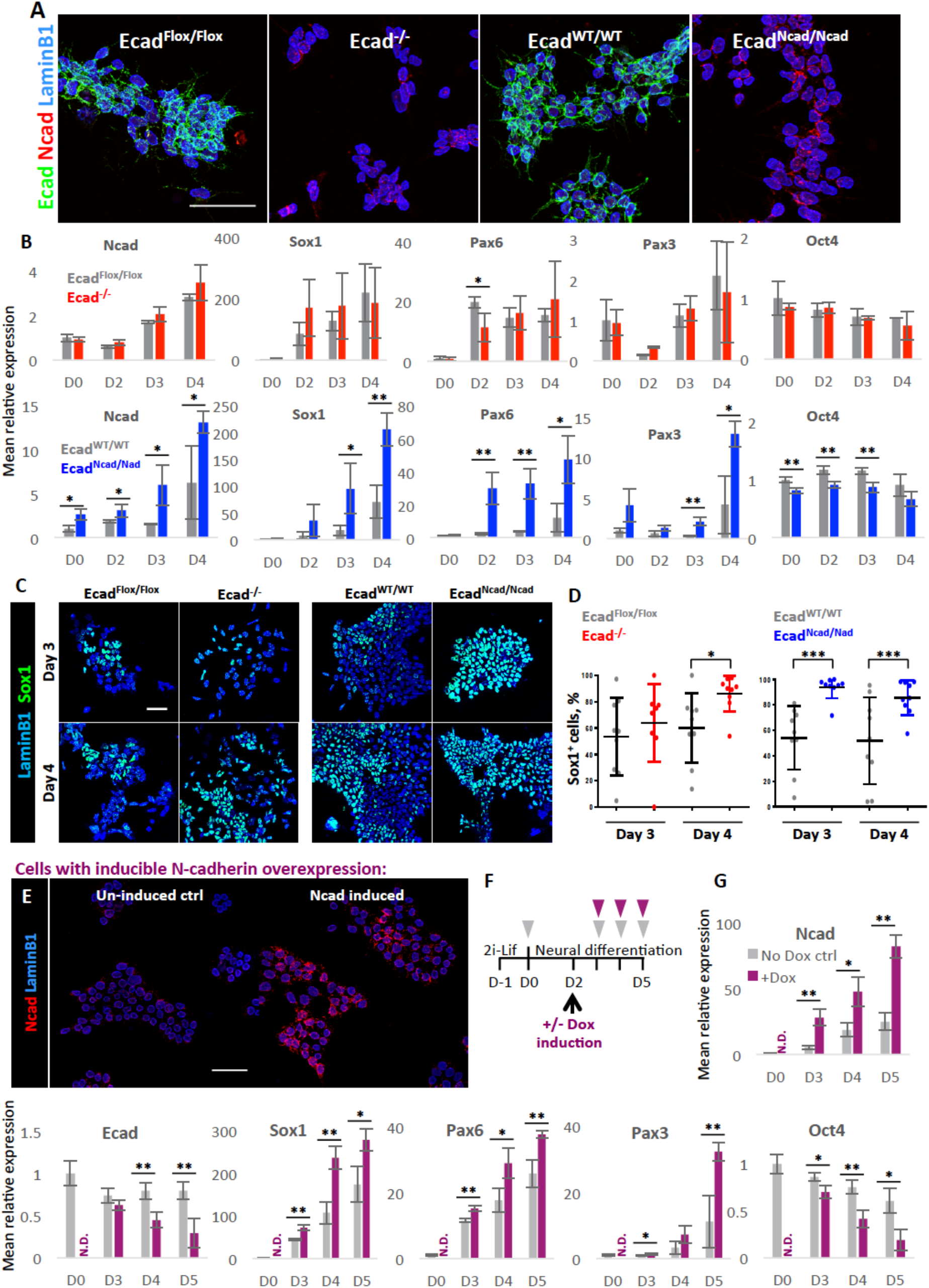
N-cadherin promotes neural differentiation. **A.** Cadherin expression in four cell lines used for cadherin domain deletion/substitution experiments. Cells cultured in neural differentiation conditions for 24h. LaminB1: nuclear envelope marker. **B.** qPCR analysis of above cell lines during successive days of neural differentiation. N=3. Values normalised to control cell line on D0. **C.** Representative images showing expression of the early neural marker Sox1 in above cells after 3-4 days of neural differentiation. **D.** Quantification of Sox1 expression in the above cells during neural differentiation. N=9, three fields of view from three biological replicates. **E.** ICC of inducible N-cadherin overexpressing cells cultured in neural differentiation conditions for 24h with or without Dox. **F.** Protocol for neural differentiation of inducible N-cadherin overexpressing cells. Triangles indicate sample collection. **G.** qPCR analysis of inducible N-cadherin overexpressing cells during neural differentiation; Dox added on day 2. N=9 (three biological replicates of three independent clones). Error bars=SD, *p≤0.05, **p≤0.01, unpaired T-test. All scale bars=50μm.

Both Ecad^-/-^ and Ecad^Ncad/Ncad^ cells can be maintained in 2i-Lif conditions. When challenged with neural differentiation conditions, both cell lines were able to switch on the neural marker genes Sox1, Pax6 and Pax3 (Figs. 2B-D). Ecad^-/-^ cells displayed a moderate pro-neural phenotype with some variability in neural gene expression. However, they showed greater instability than control cells, with very few cells surviving past d4 of neural differentiation (Fig. S2). In contrast, Ecad^Ncad/Ncad^ cells exhibited a more pronounced pro-neural phenotype, differentiating more rapidly than control cells (Fig 2B-D). Quantification of Sox1 expression in individual cells indicated that while Sox1 shows considerable cell-cell variability in both control and (to a lesser extent) Ecad^-/-^ populations, this variability was largely eliminated in Ecad^Ncad/Ncad^ cells, which exhibited uniformly high Sox1 expression by day 3. These results indicated that exogenous N-cadherin reinforces rather than reverses the pro-neural phenotype of E-cadherin null cells.

In order to further test whether N-cadherin contributes to promoting neural differentiation, we designed cell lines that allowed us to force expression of N-cadherin in the presence of endogenous E-cadherin. These cells, termed A2Lox-Ncad-HA cells, are engineered to enable dox-inducible expression of an N-cadherin transgene with a C-terminal HA tag (Fig. 2E).

When exogenous N-cadherin was induced in these cells during neural differentiation (Fig. 2F), an increase in expression of early neural markers Sox1, Pax6 and Pax3, and an accelerated downregulation of the pluripotency marker Oct4 was observed (Fig. 2G). E-cadherin was also downregulated more rapidly in the presence of ectopic N-cadherin, but these differences in E-cadherin did not emerge until one day after the pro-neural phenotype first became apparent. This might suggest that the loss of E-cadherin is likely to be a consequence rather than a cause of premature neural differentiation in these experiments, although we cannot exclude the possibility that changes in E-cadherin contribute to the effects on differentiation in these experiments.

Taken together, these results showed that the switch from E-cadherin to N-cadherin can promote neural differentiation under permissive conditions, and that the presence of N-cadherin contributes to this pro-neural effect.

### Pro-neural effects of N-cadherin are not explained by changes in cell cohesiveness

Since the primary function of cadherins is in cell-cell adhesion, we assessed whether the pro-neural effects of cadherin switching might be a consequence of cells becoming less cohesive, i.e. moving further apart from one another. In culture, Ecad^-/-^ cells are unable to form large, compact colonies, instead growing as small dispersed clumps of few cells (Fig 2A) (17,18). By contrast, N-cadherin knock-in (Ecad^Ncad/Ncad^) cells appear superficially indistinguishable from control cells (Fig. 2A) in keeping with the ability of N-cadherin to rescue the adhesion phenotype of E-cadherin null blastocysts (19,39). However this does not exclude the possibility that manipulation of N-cadherin expression could have subtle effects on cell cohesion that were not discernible by eye nor did it exclude the possibility that adhesion or migration defects became apparent under differentiation conditions.

We set out to measure whether our manipulations of cadherin expression resulted in changes in cell cohesiveness. We measured the inter-nuclear distances of cells (Fig. 3A) because we are able to perform nuclear segmentation with high accuracy (42), and because pluripotent and early neural cells have very scant cytoplasm, so inter-nuclear distance is a reasonable proxy for intercellular distance. This assay was designed to indirectly capture the consequences of any changes in adhesion or morphology, which could include a relaxation of cell-cell contacts and an increase in migration. These measurements were performed in cells that had been cultured in neural differentiation conditions for 24 hours, a time at which cells are starting to initiate differentiation but have not yet committed to a neural fate (43).

**Figure 3:**
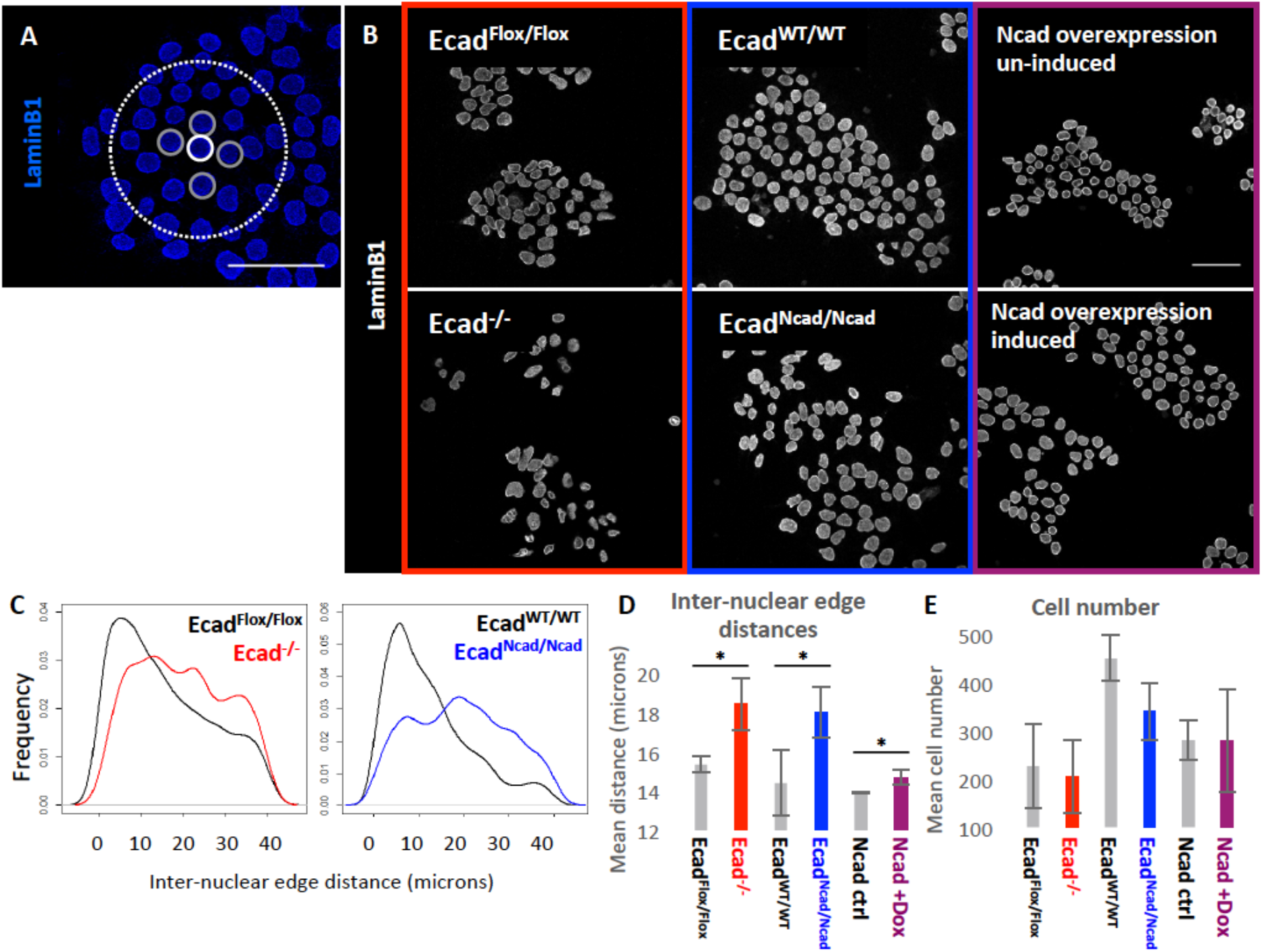
Effects of cadherin switching on cellular clustering. **A.** Methodology for measuring inter-nuclear edge distances. For each nucleus (white solid line), the nearest neighbours (grey solid line) within a 40μm radius (white dashed line) are determined by Delaunay triangulation, and the inter-nuclear edge distances between these nuclei are calculated; the process is repeated for all nuclei in the image. Scale bar=50um. **B.** ICC of all cell lines used for cadherin domain deletion/substitution experiments stained for LaminB1. Ncad overexpression was performed in A2Lox-Ncad-HA cells. Cells cultured in neural differentiation conditions for 24h, scale bar=50um. **C.** Density plots of inter-nuclear edge distance in four cell lines. N=1054 for all samples, plots show a representative sample for three biological replicates. **D.** Mean inter-nuclear edge distances. N=3 biological replicates, each containing 100-1000s of cells. **E.** Mean cell number. N=3 biological replicates. Error bars=SD, *p≤0.05, **p≤0.01, paired T-test.

We demonstrated that E-cadherin-null cells (Ecad^-/-^) had significantly greater mean inter-nuclear edge distances than control cells at 24h under differentiation conditions (Fig. 3B-D), consistent with their previously-reported adhesion defect (17,18). Surprisingly, N-cadherin knock-in cells (Ecad^Ncad/Ncad^) also had higher mean inter-nuclear edge distances from one another compared to control cells under these conditions (Fig. 3C-D). The number of cells present in each analysis did not differ significantly between the control and experimental conditions, suggesting that the observed differences in clustering were not a result of variable cell density (Fig. 3E). These results indicate that although N-cadherin can rescue the adhesion defect of E-cadherin null cells in the context of preimplantation development (39), it cannot fully rescue effects on intercellular distances (which may result from changes in adhesion, migration and/or morphology) at early stages of neural differentiation in culture. These observations therefore do not exclude the possibility that the loss of E-cadherin influences differentiation partly though changes in adhesion, migration, or morphology.

Exogenous N-cadherin can promote neural differentiation even in the presence of E-cadherin (Fig 2G), and so we next asked whether these pro-neural effects of N-cadherin correlate with a change in inter-nuclear distances. When we induced N-cadherin in the presence of endogenous E-cadherin, inter-nuclear distances were barely affected (Fig. 3D). This alerted us to the possibility that the pro-neural effect of N-cadherin might not be fully explained by changes in cell cohesiveness, and prompted us to explore whether other mechanisms may operate.

### Loss of E-cadherin leads to the loss of global and nuclear β-catenin, but does not abolish WNT responsiveness

If N-cadherin does not promote differentiation entirely through changes in cell cohesiveness, could it also be modulating pro-neural or anti-neural signalling pathways? This seems plausible since cadherins can bind various cell-surface signalling receptors and modulate signalling in other contexts (19,23,44).

We used a reverse phase protein array (RPPA) to measure the activity of a panel of signalling pathways (Supplemental Table S2) and ask which of them are changed in response to cadherin switching. We assayed signalling pathway activity by measuring changes in the abundance of total protein levels and various phospho-protein species in E-cadherin null (Ecad^-/-^) cells and N-cadherin knock-in (Ecad^Ncad/Ncad^) NcKI cells compared to their respective control cell lines. All cell lines were cultured in neural differentiation conditions for 24 hours, a time prior to the emergence of the earliest neural cells that appear in response to either endogenous or experimentally-induced cadherin switching.

We first confirmed that the most significant change in protein abundance in Ecad^-/-^ and Ecad^Ncad/Ncad^ cells was a loss of E-cadherin, as expected. The next most significant change was a depletion of β-catenin, whose levels were reduced by 3-fold both in Ecad^-/-^ and Ecad^Ncad/Ncad^ cells compared to control cells (Fig. 4A). This is in keeping with reports that β-catenin expression is reduced in response to a reduction in E-cadherin in other contexts (45,46). Ecad^-/-^ cells also have moderately reduced levels of PKA, AktP-Ser473 and AktP-Thr308, but in all cases levels of these proteins were restored to at least normal levels in Ecad^Ncad/Ncad^ cells, making it unlikely that the pro-neural phenotype could be attributed to these changes in this context. We therefore focused on β-catenin as a candidate for mediating the pro-neural effects of cadherin switching.

**Figure 4:**
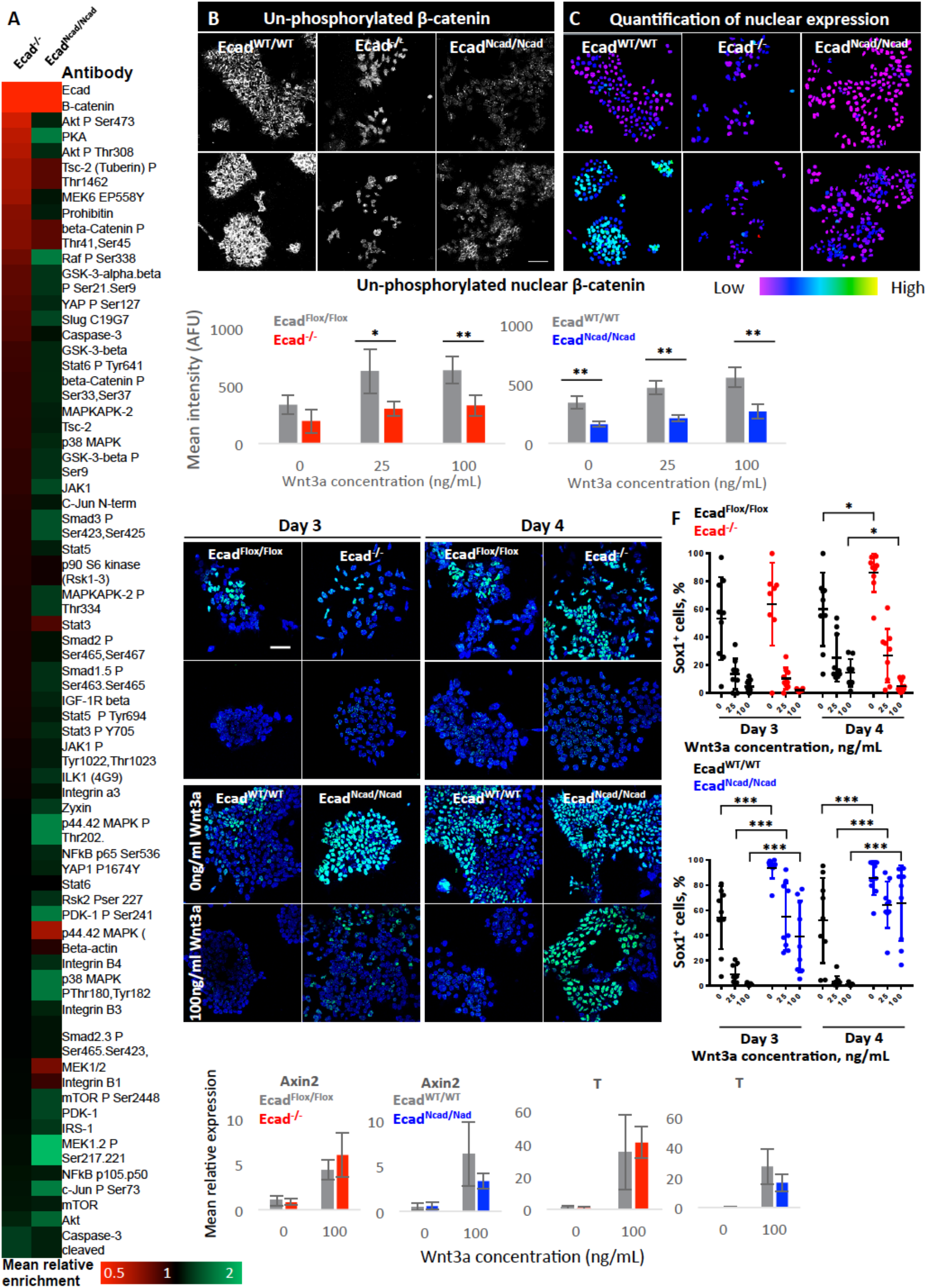
Effects of cadherin switching on β-catenin and WNT signalling. A. Heatmap showing enrichment of protein and phospho-protein species in Ecad^-/-^ and Ecad^Ncad/Ncad^ cells compared to control cell lines; all cells were cultured in neural differentiation conditions for 24h at time of analysis. Data generated by RPPA, N=3. B. Example confocal images of cells at 24h of neural differentiation cultured in varying Wnt3a concentration and stained for un-phosphorylated β-catenin. Scale bar=50μm. C. Quantitative visualisations of nuclear staining of images shown in (B). D. Quantification of mean nuclear voxel intensity for un-phosphorylated β-catenin in cells cultured for 24h in neural differentiation conditions. N=4 biological replicates, each containing 100s of cells. E, F. Example images (E) of Sox1 expression in cells cultured with or without Wnt3a. Dot plots (F) show quantification of percentage of Sox1-positive cells; each dot represents one field of view; N=9, three images each sampled from three biological replicates. G. qPCR analysis of two Wnt pathway readouts in Ecad^-/-^ and Ecad^Ncad/Ncad^ cells during neural differentiation in increasing concentrations of Wnt3a. N=3 biological replicates. For all graphs, error bars=SD, *p≤0.05, **p≤0.01, ***p ≤ 0.001; unpaired T-test.

Since β-catenin is a central player in the canonical WNT signalling pathway (47), and WNT is an anti-neural signal (27,28), we assessed whether the reduced levels of β-catenin caused changes in WNT signalling responsiveness during differentiation. We first measured nuclear accumulation of an unphosphorylated (transcriptionally active) form of β-catenin in response to Wnt3 (47). Ecad^-/-^, Ecad^Ncad/Ncad^, and control cells were cultured for 24h in serum free media with varying concentrations of Wnt3a, and the amount of un-phosphorylated β-catenin staining in the nucleus was then quantified. As expected, nuclear β-catenin increased in control cells in response to increasing concentrations of Wnt3a (Figs. 4B-D). Strikingly, Ecad^-/-^ and Ecad^Ncad/Ncad^ cells accumulated significantly less nuclear β-catenin compared to WT cells even at the high dose of Wnt3a. These results show that E-cadherin null cells display a dampened response to Wnt3a at least at the level of β-catenin accumulation and that this is not rescued by N-cadherin.

We next asked whether cadherin switching allowed cells to resist the anti-neural effects of Wnt signalling. We first confirmed that the addition of Wnt3a suppressed neural differentiation in both wild type cells and Ecad^-/-^ cells, as previously reported (27,28)(Fig 4E, F). In contrast, Ecad^Ncad/Ncad^ cells upregulated early neural markers even in the presence of high concentrations of Wnt3a, in keeping with the hypothesis that cadherin switching maintains neural potency by dampening Wnt signalling (Fig. 4E,F, Fig. S3).

However, to our surprise, WNT target genes Axin2 and T/Brachyury responded to Wnt3a in Ecad^Ncad/Ncad^ cells to a similar extent as control cell lines (Fig. 4G). This indicated that despite differences in global and nuclear levels of β-catenin, cells lacking E-cadherin were able to activate Wnt target genes normally during neural differentiation, as has been reported to be the case in other contexts (45). We speculate that this could be explained if the reduced levels of nuclear β-catenin remain above the threshold required for an efficient transcriptional response.

It is particularly interesting that Ecad^Ncad/Ncad^ cells activated the neural marker Sox1 even in the presence of the anti-neural signal Wnt3a (Figs. 4E,F): this observation suggest that cadherins become particularly important for reinforcing differentiation in a suboptimal signalling environment. However, this pro-neural effect of cadherin switching did not seem to be explained by a dampening of Wnt-responsiveness because the transcriptional response to Wnt remains intact.

### Cadherin switching dampens FGF signalling during neural differentiation

We next set out to ask which other signalling pathways are influenced by cadherin switching. Our RPPA assay indicated that a large number of proteins involved in signalling pathways were modulated in response to the concerted loss of E-cadherin and gain of N-cadherin (Fig. 4A). We previously established that exogenous N-cadherin reinforces neural differentiation (Fig 2D, Fig 4F), and so in order to simplify our search we decided to focus on signalling pathways than are modulated by N-cadherin.

We used a Nanostring assay to focus on transcriptional readouts of a broad range of signalling pathways. We measured changes in signalling pathway activity 48h after inducing N-cadherin expression during neural differentiation using our dox-inducible N-cadherin cell line (A2Lox-Ncad-HA) as this was the time-point where the pro-neural phenotype became clearly apparent.

Of 770 genes assayed (Supp Table S4), only a small number were upregulated in response to N-cadherin. These include markers of early neuroepithelial cells (Hes5: upregulated 2.4 fold, Pax3: upregulated 2.3 fold, Jag1: upregulated 1.5 fold,) consistent with a pro-neural effect of N-cadherin (Table 1). In contrast, a much larger number of genes (129 out of 142 genes at 48h post-induction) were significantly downregulated in response to N-cadherin overexpression. This suggests that N-cadherin generally suppresses rather than activates transcriptional responses to signalling pathways in this context.

**Table 1:**
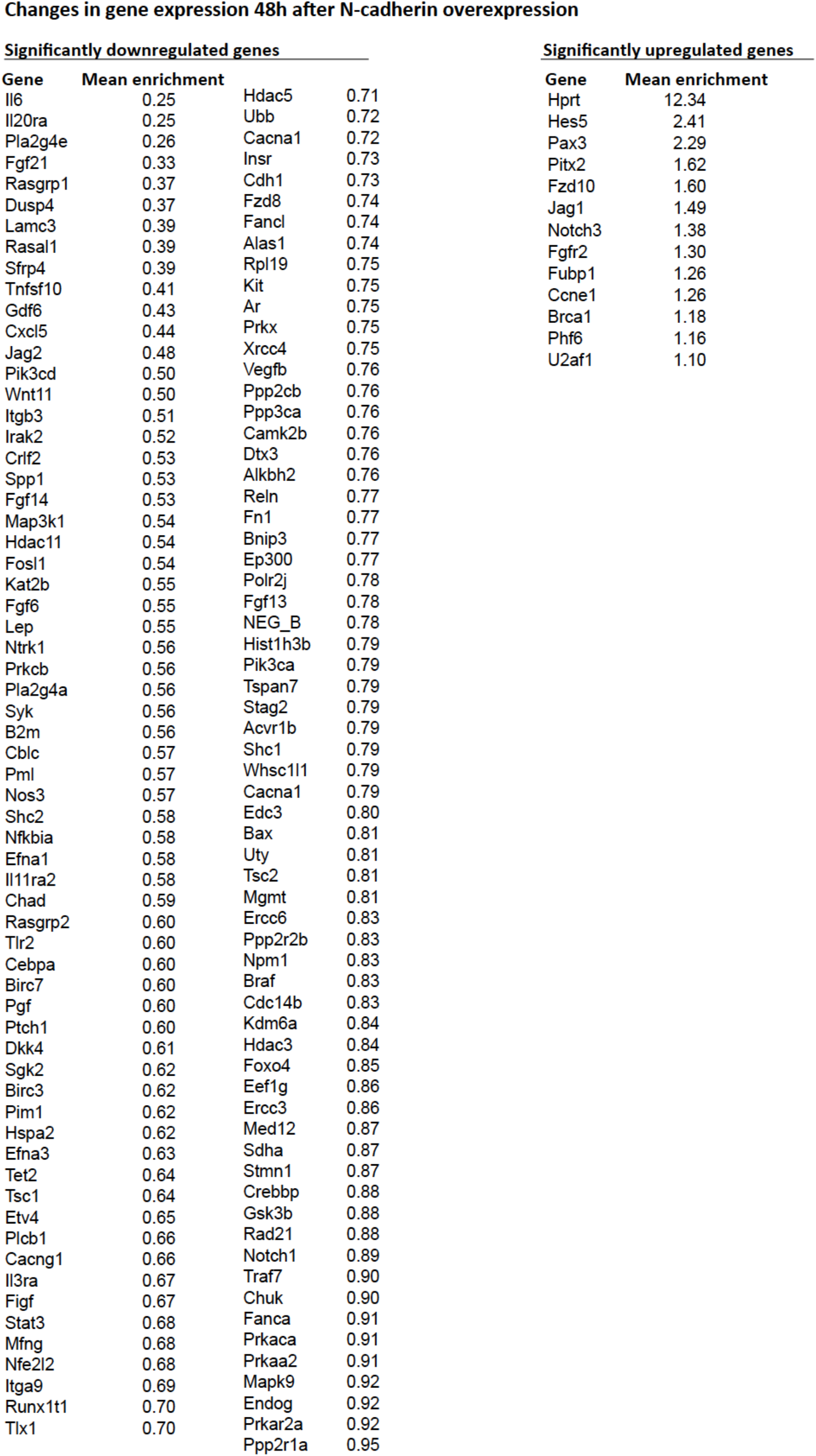
Significant changes in gene expression 48h after N-cadherin overexpression during neural differentiation. Cells were cultured for 48h in neural differentiation conditions when N-cadherin overexpression was induced by addition of Dox. RNA samples were collected for Nanostring gene expression analysis 48h later. The analysis included 770 genes involves in cellular signalling pathways. Values show mean enrichment compared to un-induced controls. N=3 biological replicates.

We then assigned each of these transcriptional changes to particular signalling pathways. This revealed that the top three pathways modulated by N-cadherin are PI3K/Akt, Ras and MAPK: all of these are pathways downstream of FGF receptors (Fig. 5A). These results indicated that N-cadherin dampens signalling pathways downstream of FGF during the early stages of neural differentiation, in keeping with reports that N-cadherin can interact with the FGF receptor in other contexts (Nguyen et al., 2016, Greber et al., 2010, Williams 1994 et al., Williams et al., 2001).

**Figure 5:**
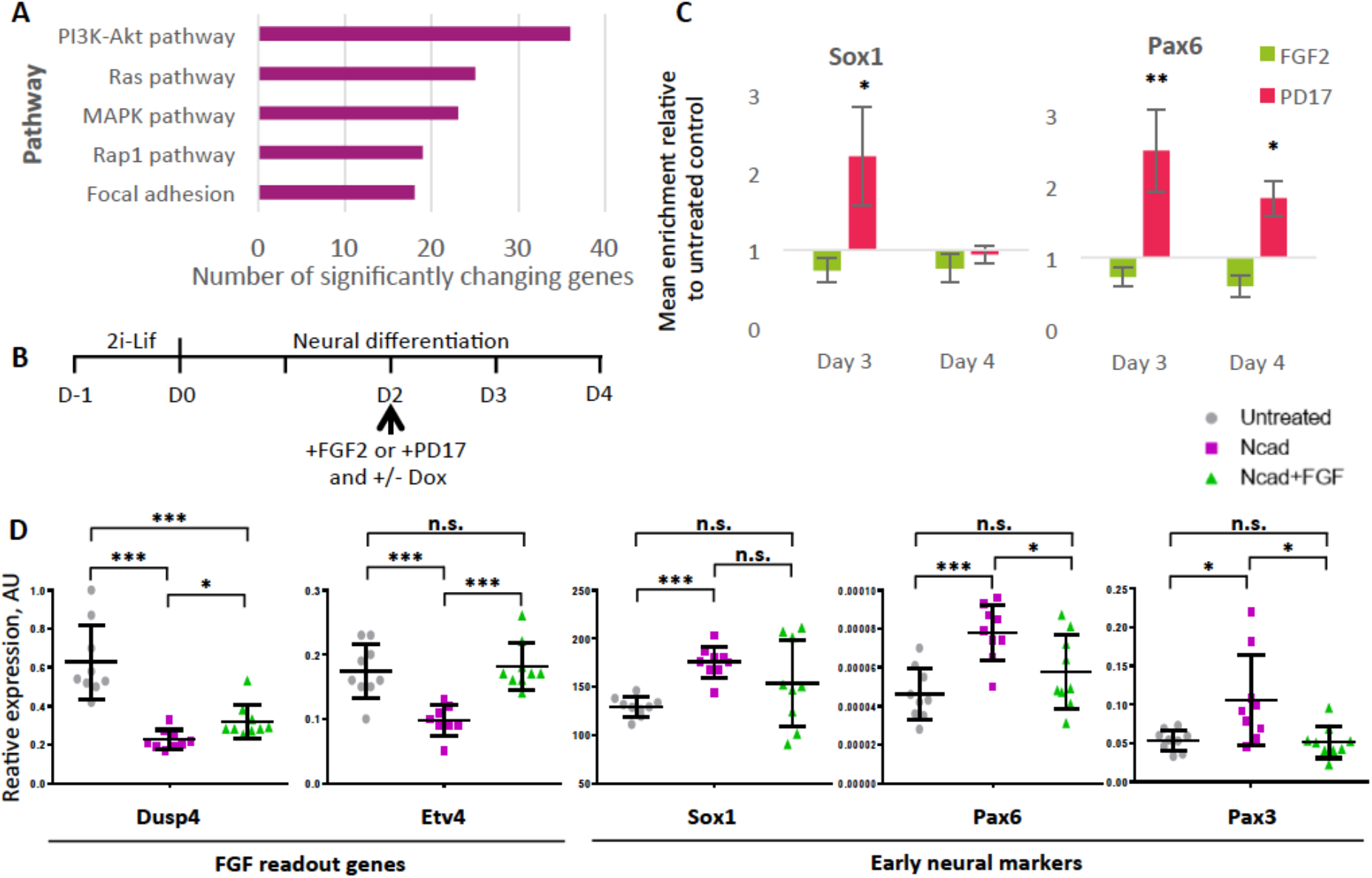
Cadherin switching promotes neural differentiation by dampening FGF signalling. **A.** Top five signalling pathways most affected by N-cadherin overexpression 48 post-induction compared to un-induced control. Data generated by DAVID functional annotation of Nanostring mRNA expression values. 142 genes out of 770 genes tested changed significantly. **B.** Protocol for FGF modulation experiments in inducible N-cadherin overexpressing cell lines. **C.** qPCR analysis in WT cells with FGF2 or PD17. N=3 biological replicates, error bars=SE *p≤0.05, **p≤0.01 compared to untreated control, unpaired T-test. **D.** qPCR analysis of gene expression after 48h of N-cadherin induction. N=9, error bars=SD, *p≤0.05, **p≤0.01, unpaired T-test. N.s.= no significance.

### N-cadherin promotes neural differentiation by dampening FGF signalling

We next assessed whether dampening of FGF signalling explains the pro-neural effect of cadherin switching.

It has previously been reported that FGF signalling promotes the acquisition and maintenance of primed pluripotency, but must then be downregulated in order for primed cells to progress to a neural fate (23–26). In keeping with these reports, we found that blockade of the FGFR1 receptor using 100ng/mL of the pharmacological inhibitor PD173074 enhances the efficiency of neural differentiation when added at d2. Conversely, addition of 20ng/mL of FGF2 reduces expression of the early neural markers Sox1 Pax6 when added at d2 (Figs. 5B, C).

Having confirmed that FGF can act as an anti-neural signal in this context, we set out to test the hypothesis that N-cadherin promotes neural differentiation by dampening FGF responsiveness. In order to test this idea we asked whether boosting FGF activity was able to reverse the pro-neural effect of N-cadherin.

We used dox-inducible N-cadherin (A2Lox-Ncad-HA) cells in order to induce N-cadherin over-expression during neural differentiation. FGF2 or the FGFR1 inhibitor PD173074 were added in order to modulate FGF activity. We added these reagents at the same time that dox was added to induce N-cadherin (Fig. 5B). We used FGF target genes Etv4 and Dusp4 to monitor FGF activity and demonstrated that N-cadherin dampens FGF activity during neural differentiation. We also observed that this effect could be at least partially rescued by addition of FGF2 (Fig. 5D). We then monitored neural differentiation by measuring expression of Sox1, Pax6 and Pax3. We found that addition of FGF2 reversed the pro-neural effects of N-cadherin, restoring the expression of these genes to similar levels as those seen in control cells that were undergoing differentiation in the absence of exogenous FGF or N-cadherin (Fig. 5D).

We conclude that N-cadherin can promote neural differentiation by dampening FGF signalling.

### Cadherin switching and neural differentiation are more synchronous *in vivo* than *in vitro*

N-cadherin is first detectable in a subset of undifferentiated epiblast stem cells prior to neural differentiation (Fig. 1). Cadherin switching then proceeds progressively over several days and is not completed in all cells until after neural fate is established. We next asked whether cadherin switching also occurs asynchronously over several days during neural development *in vivo*.

E-cadherin is expressed throughout the anterior epiblast during gastrulation. Although we were able to detect N-cadherin protein in EpiSC, we were unable to detect this adhesion molecule in the epiblast of mouse embryos during gastrulation: it is detected exclusively in the emerging mesoderm, as previously reported (48)(Fig. S4).

We then focused on the newly-formed Sox1^+^ cells within the neural plate. We observed that this region uniformly displays E-cadherin but lacks detectable N-cadherin until after gastrulation (Fig. 6A, Fig. S4). In contrast to the heterogeneity in cadherin expression observed before and during neural differentiation in culture (Figs. 1A, 1H), we were unable to detect any obvious local cell-to-cell variability in expression of either E- or N-cadherin within the epiblast or the early neuroepithelium, (Fig. 6A). We also noticed that almost every cell expresses Sox1 in the anterior neuroepithelium *in vivo*, while in contrast only around a third of cells express Sox1 during an equivalent stage of neural differentiation in culture (Figs. 6B-F).

**Figure 6:**
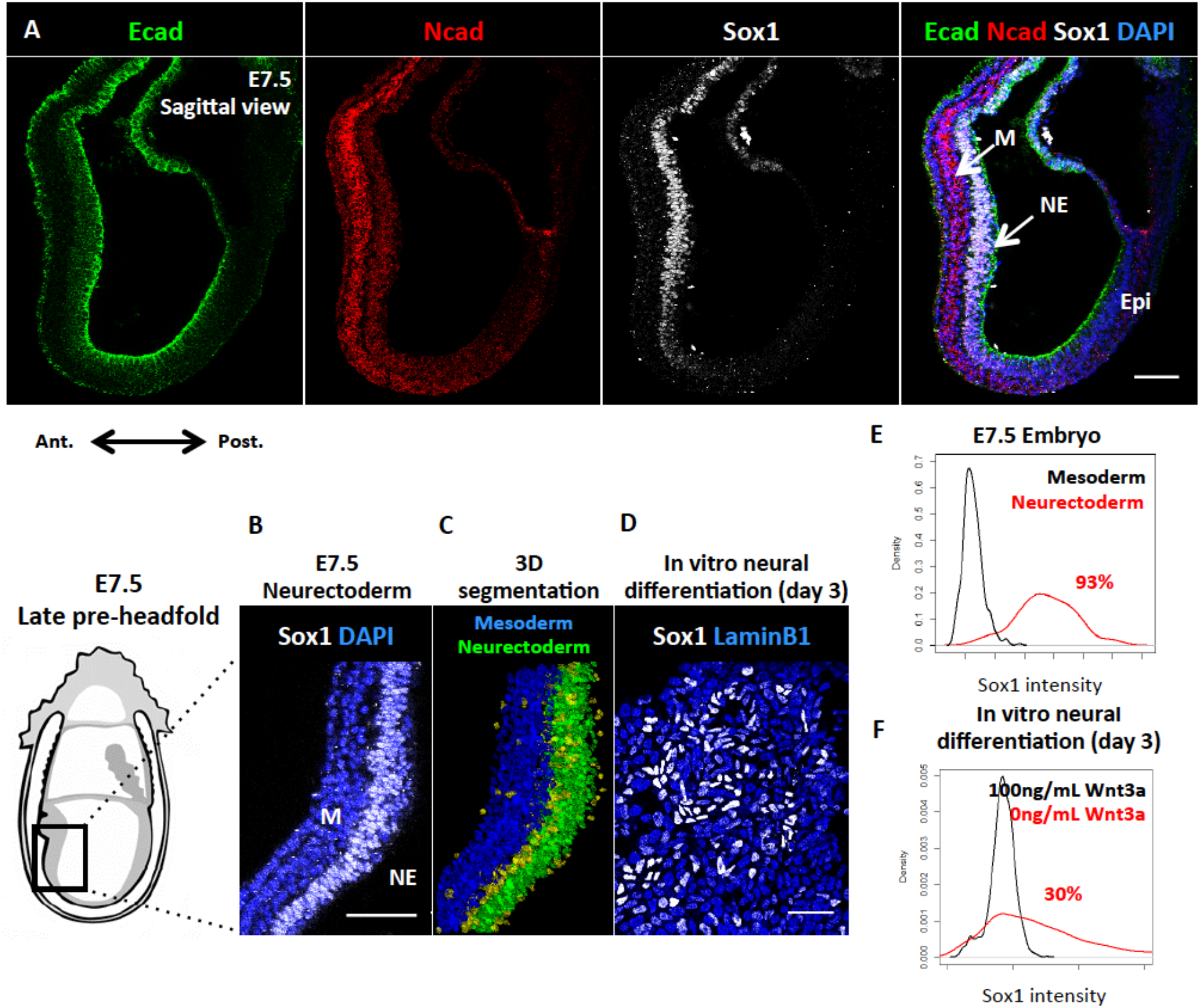
Cadherin switching and neural differentiation are more synchronous *in vivo* than *in vitro*. **A.** E- and N-cadherin co-expression in the E7.5 mouse embryo, sagittal view. M: mesoderm; N: neuroectoderm; Epi: epiblast. **B.** Sox1 expression in the anterior neurectoderm at E7.5. Nuclei stained with DAPI. **C.** Nuclear segmentation and binning of cells shown in (B). **D.** Sox1 expression at day 3 of neural differentiation in vitro. Nuclei stained with nuclear envelope marker LaminB1. **E.** Quantitative analysis of Sox1 expression in two tissues of the anterior E7.5 embryo. N=438 nuclei per sample. **F.** Quantitative analysis of Sox1 expression at day 3 of neural differentiation in vitro, cultured with and without the neural differentiation inhibitor Wnt3a. N=231 nuclei per sample. In F-G, percentages refer to the proportion of Sox1-positive cells in the neural sample (red line; negative control population in black) as calculated by the non-overlapping area under the two curves. Embryo drawings are adapted from the EMAP eMouse Atlas Project (http://www.emouseatlas.org). All scale bars=50μm.

We conclude that in the embryo, cells acquire N-cadherin after acquisition of neural identity, consistent with a role in stabilising rather than inducing neural fate. We also observe that cadherin switching occurs relatively synchronously *in vivo*, in contrast with the asynchronous acquisition of N-cadherin in culture. Given our finding that N-cadherin can stabilise neural identity, this dysregulation of cadherin switching in vitro may help to explain why neural differentiation proceeds less synchronously in culture than in the embryo.

## Discussion

Here we report that the switch from E-to N-cadherin helps to reinforce neural commitment by dampening FGF signalling. It has previously been reported that premature cadherin switching results in gross morphological and cell-fate allocation defects at gastrulation, resulting at least in part from defects in extra-embryonic tissues (5). Our findings suggest that there may also be a cell-autonomous requirement for cadherin switching during neural differentiation.

E-cadherin is required to initiate differentiation in some contexts (40), but once differentiation is triggered cadherins can have positive or negative effects on subsequent lineage specification (32,40), highlighting the multiple stage-specific effects of cadherins during differentiation of pluripotent cells. Our findings confirm previous reports that the absence of E-cadherin can limit the pool of nuclear beta catenin (45,46,49), but we find that this does not result in a dampening the transcriptional response to Wnt in differentiating neural progenitors: this is in keeping with similar findings in some cell types (45,46,49), but contrasts with findings in other contexts where changes in E-cadherin do modulate the transcriptional response to Wnt (33,41). Nevertheless, cadherin switching enables cells to resist the anti-neural effects of Wnt, possibly through an indirect mechanism. It would be interesting to explore the positional identity and potency of the Sox1^+^ cells that emerge in the presence of exogenous Wnt and N-cadherin, given that Sox1 is expressed in neuromesodermal progenitors (50) and that Wnt helps support neuromesodermal progenitor identity in the posterior of the embryo (51,52).

We find that N-cadherin can dampen FGF activity, leading us to speculate that N-cadherin might contribute to reinforcing neural commitment by protecting early neural cells from fluctuations in the anti-neural pro-mesoderm FGF signal. Because N-cadherin on one cell will stabilise N-cadherin on neighbouring cells through homotypic interaction, it is tempting to speculate that N-cadherin helps to propagate this neural-stabilisation effect through the tissue via a type of ‘community effect’. This could help ensure that neural commitment proceeds robustly in the embryo. It has recently been proposed that cadherins propagate mesodermal differentiation from cell to cell in an *in vitro* model of the primitive streak, although in that case communication is propagated predominantly through changes in E-cadherin rather than N-cadherin (53). Differentiation is more variable and unpredictable in culture compared to the embryo, even though the extrinsic signalling environment in a culture dish can be tightly controlled. We speculate that a cadherin-mediated community effect may operate less efficiently in culture where the earliest N-cadherin positive cells will often encounter neighbours that lack N-cadherin.

This work highlights the importance of changes in adhesion and morphology in ensuring robust development, and suggests that efforts to mimic these changes in culture will be critical for gaining full control over differentiation of pluripotent cells.

## Acknowledgments

We are grateful to the staff of the MRC Centre for Regenerative Medicine Tissue Culture facility, FACS facility, and Imaging Facility for expert technical assistance. Embryo drawings in Figs 6 and S4 are adapted from the EMAP eMouse Atlas Project (http://www.emouseatlas.org)(54). We thank Lesley Forrester and Val Wilson for advice and comments on the manuscript. This work was funded by Wellcome Trust Senior Fellowship WT103789AIA to SL, by Sir Henry Wellcome Fellowship WT100133 to GB and by an MRC PhD studentship to KP.

## Materials & Methods

### Mouse ES cell culture

**Naïve stem cell culture (2i-Lif):** naïve ES cells were maintained in 2i-Lif medium (6). This medium is N2B27 supplemented with 1μM PD0325901, 3μM Chiron 99021 and 100 units/mL LIF on laminin-coated tissue culture plates. Naïve stem cells were derived by passaging LIF-serum-cultured ES cells into 2i-Lif conditions, and maintaining them for at least three passages. **LIF-serum culture:** ES cells were maintained in GMEM supplemented with 2-mercaptoethanol, non-essential amino acids, glutamine, pyruvate, 10% foetal calf serum (FCS) and 100 units/mL LIF on gelatinised tissue culture flasks (55). **Epiblast stem cell culture:** EpiSCs were maintained under in published conditions (8,9). Briefly, the cells were maintained in N2B27 solution supplemented with 20ng/mL Activin and 10ng/mL FGF on fibronectin-coated tissue culture plates. EpiSCs were derived by passaging LIF-serum-cultured ES cells into EpiSC conditions, and maintaining them for at least three passages.

### Cell lines

E14tg2α mouse embryonic stem cells were used as WT cells. Previously published cell lines were generously provided by the researchers and labs who generated them: Sox1-GFP (“46C”) cells, (56); Ecad^Ncad/Ncad^ cells (5,39) that were targeted to an E14.1 background, and chimeric mice were then backcrossed to C57BL/6 for at least 3-4 generations before homozygous Ecad^Ncad/Ncad^ ES cells were established from blastocysts; Ecad^-/-^ cells on an E14-IB10 background, in which exons 4 to 15 were Cre-excised *in vitro* (40,57).

To generate Dox-inducible N-cadherin overexpressing ES cells, the inducible cassette exchange (ICE) method was used (58,59). Primers were designed to allow for the amplification of a DNA fragment containing the *Cdh2* gene C-terminally tagged with an influenza virus hemagglutinin (HA) tag; the whole construct was flanked by XhoI and NotI restriction sites. The construct was ligated into a pCR Blunt II Topo vector (Thermo Fisher Scientific). The *Cdh2-HA* insert was then ligated into a p2Lox-eGFP plasmid, replacing an *eGFP* sequence in this construct (60). The resulting p2Lox-Cdh2-HA plasmid was then nucleofected into A2LoxCre cells. The cells were then cultured for 10 days under G418 selection, and surviving clonal colonies were then expanded. Clones were then screened for Ncad and HA expression by ICC, and three clones with high transgene expression were selected for use in further experiments.

### Differentiation protocols

Monolayer neural differentiation was performed by passaging 2i-Lif-cultured ES cells at low density into laminin-coated tissue culture plates. The cells were maintained in 2i-Lif for 24h to allow the cells to properly adhere to the matrix. After 24h, media were changed to N2B27 medium in which commercial N2 was replaced with 0.5% modified N2 (made in-house as described in Pollard et al., 2006). Media were changed every 1-2 days.

### qRT-PCR

Primers used for qRT-PCR are described in Supplementary table S1. All expression values were normalised to the geometric mean expression value of at least two of three housekeeping genes: *TBP, SDHA*, and *Ywhaz*.

### RPPA analysis

RPPA analysis was performed on nitrocellulose coated slides as previously described (62). Briefly, cells were washed with PBS and lysed in 1% Triton X-100, 50 mM HEPES (pH 7.4), 150 mM sodium chloride, 1.5 mM magnesium chloride, 1 mM EGTA, 100 mM sodium fluoride, 10 mM sodium pyrophosphate, 1 mM sodium vanadate, 10% glycerol, supplemented with cOmplete ULTRA protease inhibitor and PhosSTOP phosphatase inhibitor cocktails (Roche). Cleared lysates were serially diluted to produce a four step doubling dilution series of each sample, which were spotted in technical triplicate onto nitrocellulose-coated slides (Grace Bio-Labs) under conditions of constant 70% humidity using an Aushon 2470 array platform (Aushon Biosystems). After hydration slides were blocked using SuperBlock (TBS) blocking buffer (Thermo Fisher Scientific) and incubated with validated primary antibodies (all diluted 1:250 in SuperBlock; Supplementary table S2). Bound antibodies were detected by incubation with anti-rabbit DyLight 800-conjugated secondary antibody (New England BioLabs). Slides were analysed using an InnoScan 710-IR scanner (Innopsys), and images were acquired at the highest gain without saturation of the fluorescence signal. The relative fluorescence intensity of each array feature was quantified using Mapix software (Innopsys).

Primary antibodies used in this assay are listed in Supp table S2. The linear fit of the dilution series of each sample was determined for each primary antibody, from which median relative fluorescence intensities (RFI) values were calculated to provide relative quantification of total protein and phosphoprotein abundance across the sample set. Finally, signal intensities were normalized to total protein loading for each sample by using readout from a fast-green (total protein) stained array slide. Enrichment values for EcKO and NcKI cells were normalised to those in relevant control cell lines, and a mean enrichment was then calculated for three biological replicates.

### Statistical analysis

All experiments that were statistically analysed were performed with at least three independent biological replicates. Statistical significance was calculated using a paired or unpaired Student’s t-test as appropriate.

### Immunofluorescence, FACS

For immunofluorescence analysis, cells cultured on glass coverslips were fixed in 4% formaldehyde and incubated for at least 30 min in blocking buffer (PBS, 3% donkey serum and 0.1% Triton). Primary antibodies were diluted in blocking buffer and applied for 1 hr at room temperature or overnight at 4°C. After three washes in PBS, secondary antibodies conjugated to Alexa fluorophores (Life Technologies) were diluted at 1:1000 in blocking buffer and applied for 1 hr at room temperature. The cells were washed at least three times and the coverslips were mounted with Prolong Gold Antifade Reagent (Life Technologies) on glass slides for viewing.

For antibody staining of live cells for flow cytometry, cells were incubated with relevant antibodies on ice in the dark for at least 15 minutes. Antibodies were diluted in FACS buffer (5% FCS in PBS). Flow cytometric analysis was performed using a BD Accuri flow cytometer. FACS was carried out on a FACS Aria cell sorter.

Embryos were fixed with 4% formaldehyde/PBS/0.1% Triton X-100 (Sigma) for 30 minutes, and quenched with 50mM ammonium chloride. Cellular permeabilization was carried out for 10 min in PBS/0.1% Triton X-100. The embryos were incubated in primary antibody in 3% donkey serum/PBS/0.1% Triton X-100 overnight, and subjected to 3 washes in PBS/0.1% Triton X-100. Secondary antibodies were applied subsequently for 2h to overnight, followed by 3 washes in PBS/0.1% Triton X-100. Embryos were then stained with DAPI (Biotium), mounted in PBS droplets covered with mineral oil in “microscope rings”, and imaged on a Leica SP8 confocal microscope.

Alternatively, following staining, chimaeric embryos requiring immunostaining quantification were dehydrated in methanol series in PBS/0.1% Triton X-100, clarified in 50% methanol/50% BABB (benzyl alcohol:benzyl benzoate 1:2 ratio, Alfa Aesar and Sigma), transferred into 100% BABB in glass capillaries and imaged on a Leica SP8 confocal microscope.

### Animal Care and Use

Animal experiments were performed under the UK Home Office project license PEEC9E359, approved by the Animal Welfare and Ethical Review Panel of the University of Edinburgh and within the conditions of the Animals (Scientific Procedures) Act 1986.

### Transcript enrichment analysis

Gene enrichment datasets generated by Nanostring were analysed using the associated NSolver 4.0 software. Functional annotation of gene lists was performed using the DAVID functional annotation online tool (63,64). A list of significantly changing genes was used as an input list, while all 770 genes tested in the Nanostring analysis (Supp Table S4) were used as a background list.

### Quantitative image analysis

**Quantification of membrane-bound protein staining:** where fluorescent signal was to be quantified at the single-cell level, cells were imaged in 3D Z-stacks on a Leica SP8 three-detector confocal microscope. For the quantification of membrane staining, cells were counted manually using Fiji/ImageJ software. **Quantification of nuclear protein staining:** for the quantification of nuclear staining, PickCells software was used. Cells were segmented by nuclear content or nuclear membrane staining (using DAPI or nuclear envelope marker LaminB1 staining, respectively) using the software’s inbuilt NESSys nuclear segmentation module (42). Protein expression was then quantified as the mean pixel intensity in any given nucleus. Where staining quality did not allow for accurate nuclear segmentation, cells were manually designated as either positive or negative based on a single empirical threshold for all images generated from a single biological replicate. **Quantification of inter-nuclear edge distances:** inter-nuclear edge distance was calculated using PickCells software in cells segmented using nuclear membrane staining. The nearest neighbours for each nucleus were determined using Delaunay triangulation, and the distance between the membranes of nearest neighbours was calculated for each nucleus up to a distance of 40 microns.

## Supplementary tables

**Supplementary table S1:**
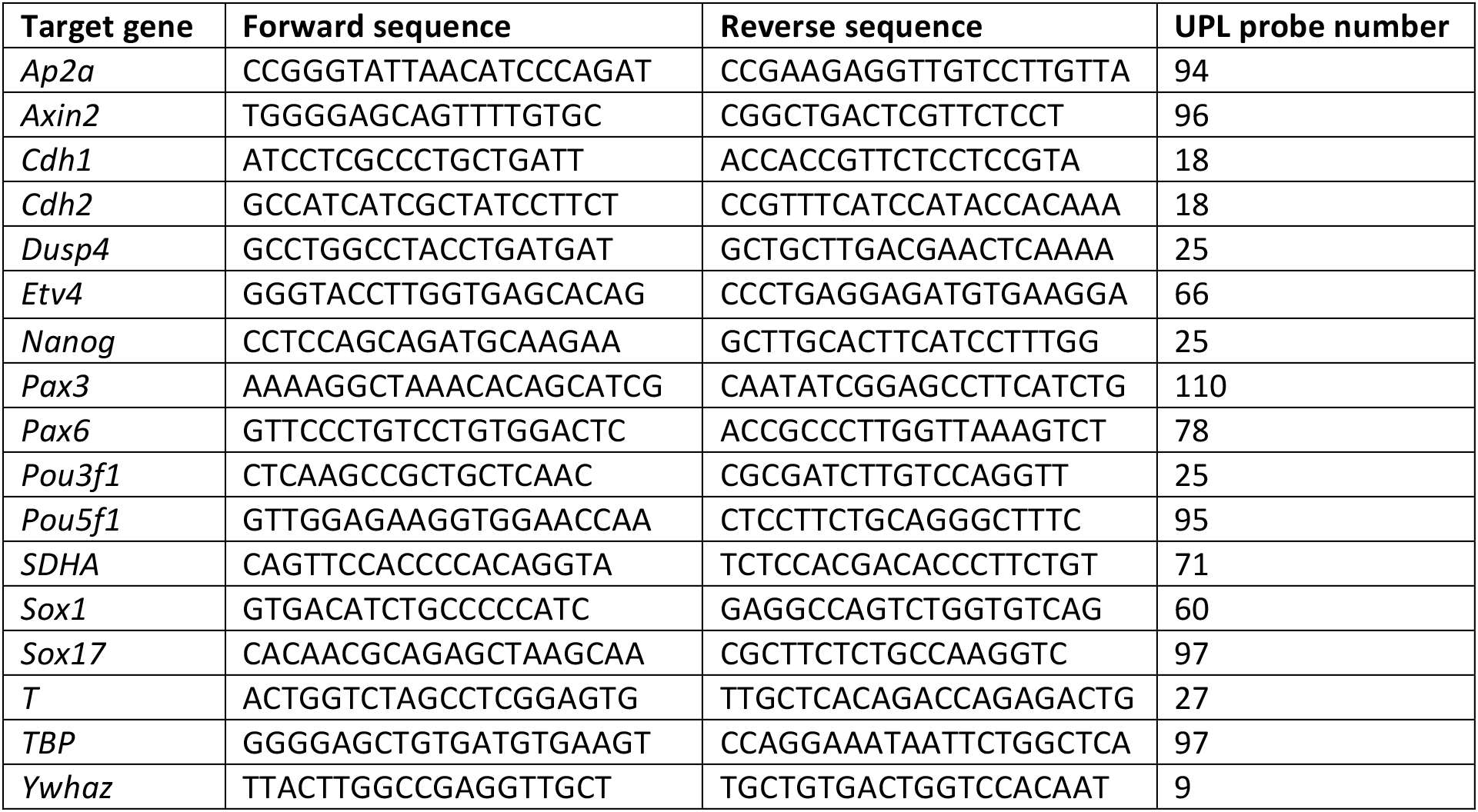
qPCR primer sequences. Primers used with the Universal Probe Library qPCR system from Roche.

**Supplementary table S2:** RPPA antibodies. All antibodies used were raised in rabbit.

| Epitope recognised | Supplier | Catalogue no. |
| --- | --- | --- |
| Akt | Cell Signaling Technologies | 9272 |
| Akt P Ser473 | Cell Signaling Technologies | 4060 |
| Akt P Thr308 | Cell Signaling Technologies | 2965 |
| beta-actin | Cell Signaling Technologies | 4970 |
| beta-Catenin | Cell Signaling Technologies | 9562 |
| beta-Catenin P Ser33,Ser37,Thr41 | Cell Signaling Technologies | 9561 |
| beta-Catenin P Thr41,Ser45 | Cell Signaling Technologies | 9565 |
| Caspase 3 | Cell Signaling Technologies | 9662 |
| Caspase 3 cleaved | Cell Signaling Technologies | 9664 |
| c-Jun N-term | Epitomics | 1254-1 |
| c-Jun P Ser73 | Cell Signaling Technologies | 9164 |
| E-Cadherin | Cell Signaling Technologies | 3195 |
| GSK-3-alpha/beta P Ser21/Ser9 | Cell Signaling Technologies | 9331 |
| GSK-3-beta | Cell Signaling Technologies | 9315 |
| GSK-3-beta P Ser9 | Cell Signaling Technologies | 9336 |
| IGF-1R beta | Cell Signaling Technologies | 3027 |
| ILK1 (4G9) | Cell Signaling Technologies | 3856 |
| Integrin alpha 4 | Cell Signaling Technologies | 4600 |
| Integrin Beta 1 [EP1041Y] | Abcam | ab52971 |
| Integrin beta3 | Cell Signaling Technologies | 4702 |
| Integrin beta4 | Cell Signaling Technologies | 4707 |
| IRS-1 | Cell Signaling Technologies | 2382 |
| JAK1 | Cell Signaling Technologies | 3332 |
| JAK1 P Tyr1022,Thr1023 | Invitrogen (Biosource) | 44-422G |
| MAPKAPK-2 | Epitomics | 1497-1 |
| MAPKAPK-2 P Thr334 | Cell Signaling Technologies | 3041 |
| MEK1/2 | Cell Signaling Technologies | 9122 |
| MEK1/2 P Ser217/221 | Cell Signaling Technologies | 9154 |
| MEK6 [EP558Y] | Abcam | ab52937 |
| mTOR | Cell Signaling Technologies | 2972 |
| mTOR P Ser2448 | Cell Signaling Technologies | 2971 |
| NFkB p105/p50 | Calbiochem | GTX110585 |
| NFkB p65 Ser536 | Cell Signaling Technologies | 3033 |
| p38 MAPK | Cell Signaling Technologies | 9212 |
| p38 MAPK PThr180,Tyr182 | Cell Signaling Technologies | 9211 |
| p44/42 MAPK (ERK1/2) | Cell Signaling Technologies | 9102 |
| p44/42 MAPK (ERK1/2) P<br>Thr202/Thr185,Tyr204/Tyr187 | Cell Signaling Technologies | 4370 |
| p90 S6 kinase (Rsk1-3) | Santa Cruz | sc-231 |
| PDK-1 | Cell Signaling Technologies | 3062 |
| PDK-1 P Ser241 | Cell Signaling Technologies | 3061 |
| PKA | Abcam | ab26322 |
| Prohibitin | Santa Cruz | sc-28259 |
| Raf P Ser338 | Cell Signaling Technologies | 9427 |
| Rsk2 Pser 227 | Cell Signaling Technologies | 3556 |
| Slug (C19G7 | Cell Signaling Technologies | 9585 |
| Smad1/5 P Ser463/Ser465 | Cell Signaling Technologies | 9516 |
| Smad2 P Ser465,Ser467 | Cell Signaling Technologies | 3108 |
| Smad2/3 P<br>Ser465/Ser423,Ser467/Ser425 | Cell Signaling Technologies | 8828 |
| Smad3 P Ser423,Ser425 | Cell Signaling Technologies | 9520 |
| Stat3 | Cell Signaling Technologies | 12640 |
| Stat3 P Y705 | Cell Signaling Technologies | 9131 |
| Stat5 | Invitrogen (Biosource) | 44-368G |
| Stat5 P Tyr694 | Cell Signaling Technologies | 9351 |
| Stat6 | Cell Signaling Technologies | 9362 |
| Stat6 P Tyr641 | Cell Signaling Technologies | 9361 |
| Tsc-2 (Tuberin) | Cell Signaling Technologies | 3612 |
| Tsc-2 (Tuberin) P Thr1462 | Cell Signaling Technologies | 3617 |
| YAP P Ser127 | Cell Signaling Technologies | 4911 |
| YAP1 [EP1674Y] | Abcam | ab52771 |
| Zyxin | Cell Signaling Technologies | 3553 |

**Supplementary table S3:** Immunocytochemistry and flow cytometry antibodies; all antibodies were diluted to the specified concentration in blocking buffer.

| Epitope recognised | Clone | Host species | Dilution factor | Supplier | Cat. number |
| --- | --- | --- | --- | --- | --- |
| β-catenin (active), dephosphorylated on Ser37 or Thr41 | 8E7 | Mouse | 1:1000 | Millipore | 05-665 |
| E-cadherin | DECMA-1 | Rat | 1:200 | Sigma | U3254 |
| E-cadherin, eFluor660-conjugated | DECMA-1 | Rat | 1:300 | eBioscience | 50-3249-82 |
| GFP | Polyclonal | Chicken | 1:1000 | Abcam | 13970 |
| HA | HA-7 | Mouse | 1:1000 | Sigma | H3663 |
| Laminβ1 | Polyclonal | Chicken | 1:1000 | Abcam | Ab90169 |
| Laminβ1 | Polyclonal | Rabbit | 1:1000 | Abcam | Ab16048 |
| N-cadherin | 32 | Mouse | 1:200 | BD | 610920 |
| N-cadherin, AlexaFluor488-conjugated | Polyclonal | Sheep | 1:50 | R&D | FAB6426G |
| Oct4 | N-19 | Goat | 1:200 | Santa Cruz | SC-8628 |
| Sox1 | N23-844 | Mouse | 1:200 | Pharmigen | 560749 |

## Supplementary Figures

**Supplementary figure S1:**
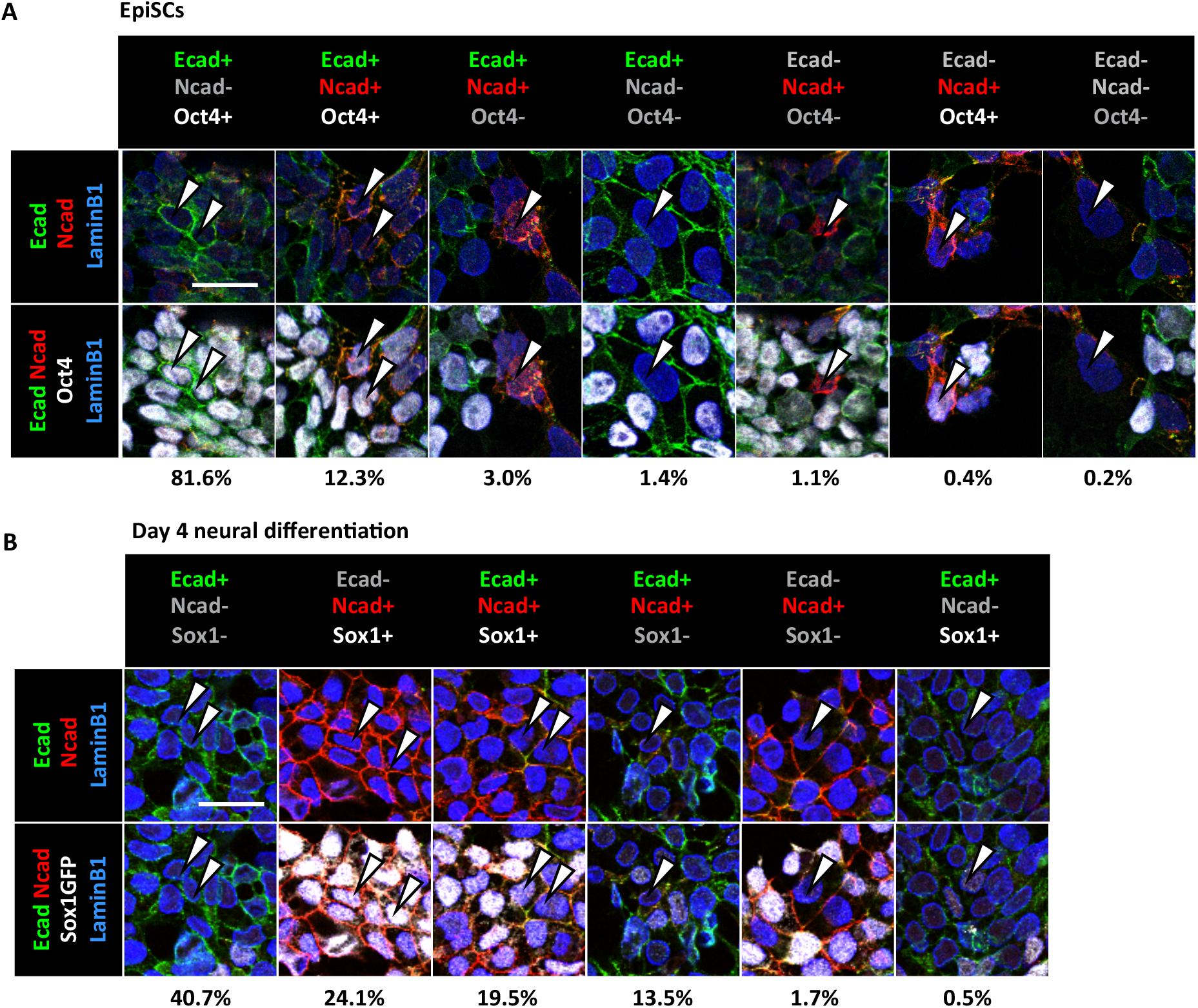
Single-cell views of protein co-expression in EpiSCs and during neural differentiation. White arrowheads point out individual cells of the given protein expression profile. Percentages below images indicate the proportion of cells of a given identity out of all cells analysed (manual quantification). Scale bars=25μm. **A.** EpiSCs stained for E-cadherin (green), N-cadherin (red), the pluripotency marker Oct4 (white) and nuclear envelope marker LaminB1 (blue). N=2596 cells from three biological replicates **B.** Cells on day four of neural differentiation from a 2i-Lif starting population stained for E-cadherin (green), N-cadherin (red), Sox1-GFP (shown in white), and nuclear envelope marker LaminB1 (blue). N=2275 cells from three biological replicates.

**Supplementary figure S2:**
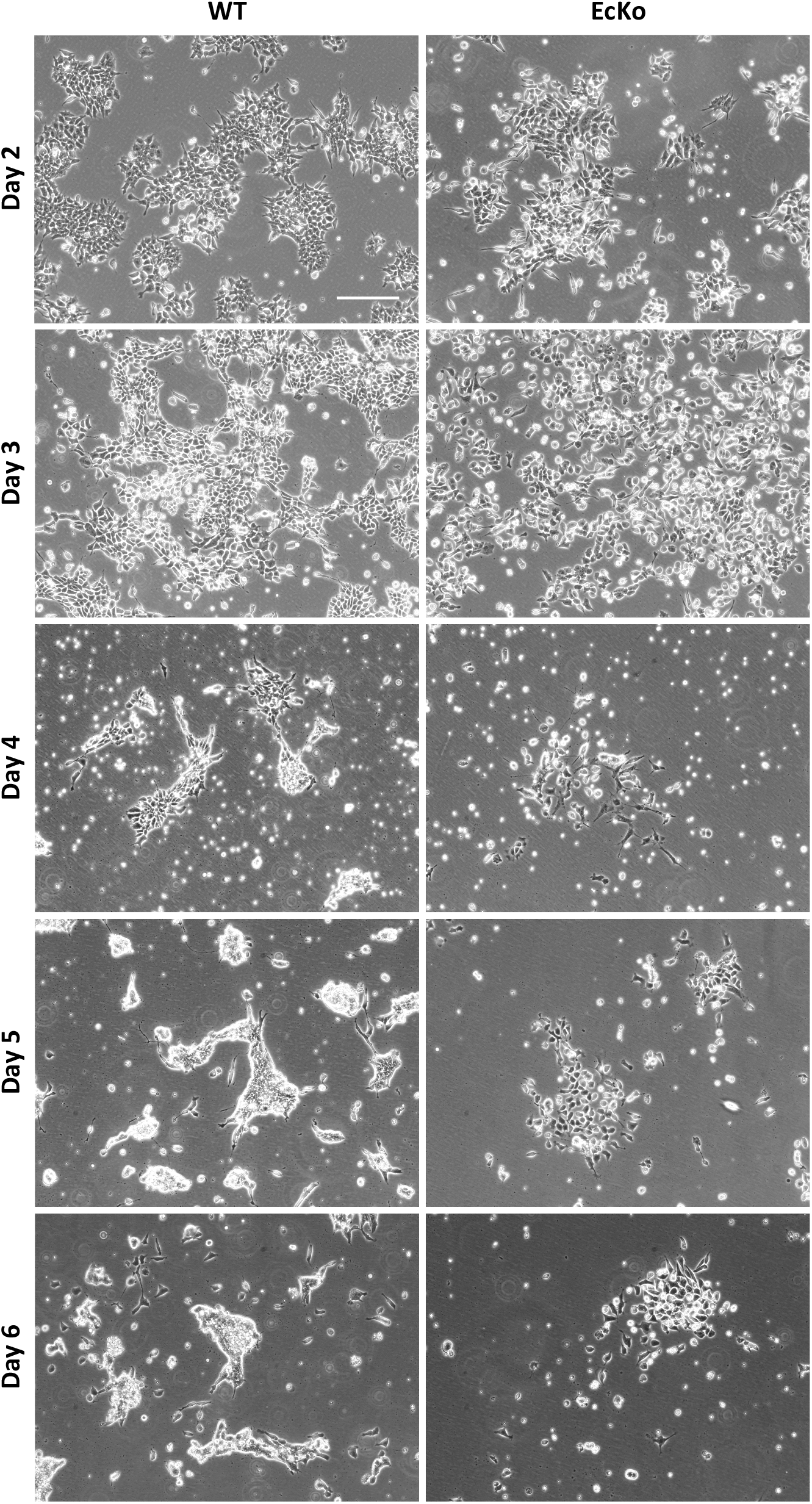
Ecad^-/-^ cells die after prolonged culture in N2B27. Phase contrast images of Ecad^Flox/Flox^ and Ecad^-/-^ cells on days 2-6 of neural differentiation from a 2i-Lif starting population. Scalebar= 100μm.

**Supplementary figure S3:**
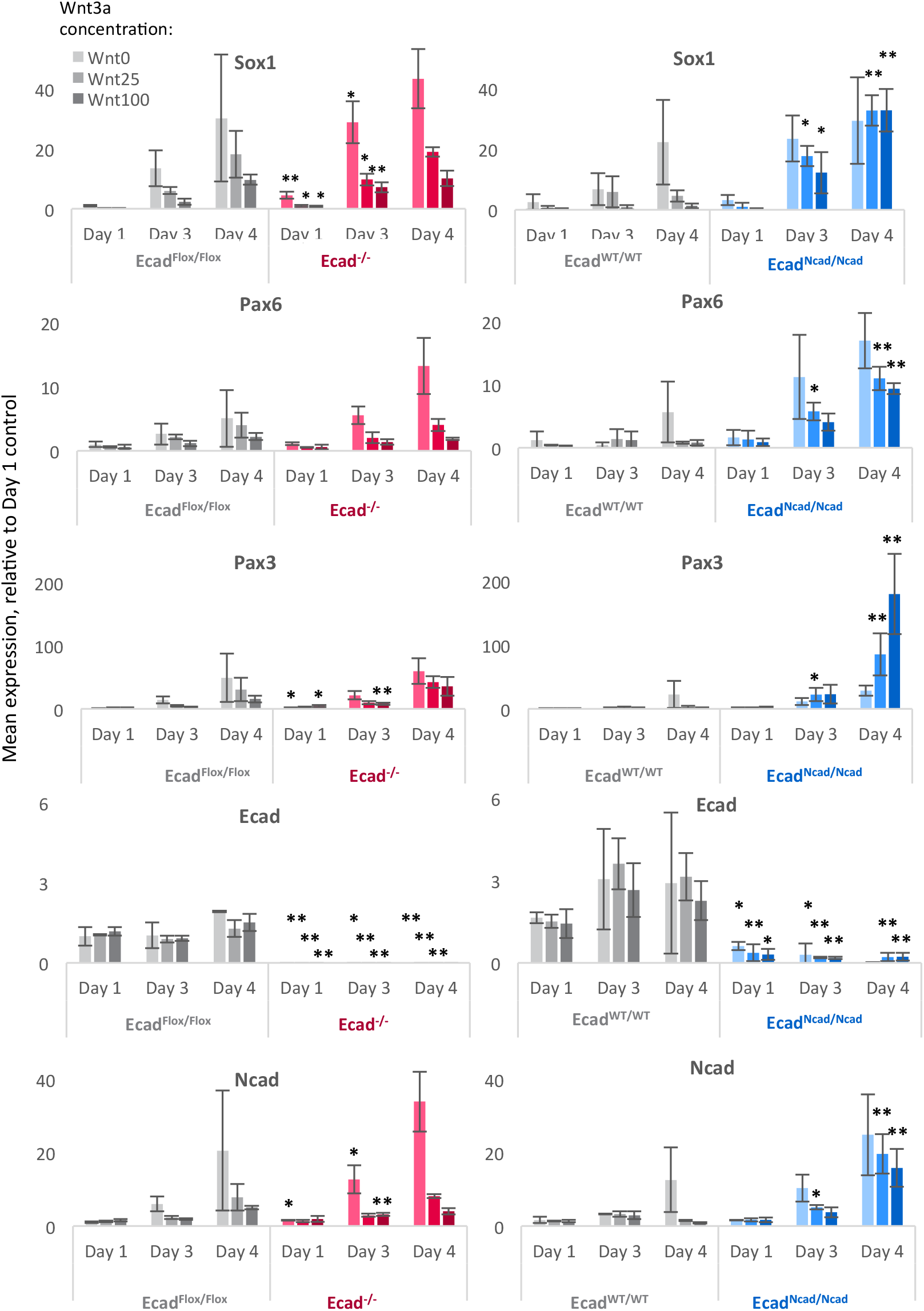
Effects of cadherin switching on β-catenin and WNT signalling. qPCR analysis of Ecad^Flox/Flox^ and Ecad^Ncad/Ncad^ cells during neural differentiation in increasing concentrations of Wnt3a; bars denote mean expression relative to the Day 1 condition of the relevant control cell line (grey). Asterisks denote significant difference compared to the paired control cell line in the same condition. N=3 biological replicates. Error bars=SD, *p≤0.05, **p≤0.01, paired T-test.

**Supplementary figure S4:**
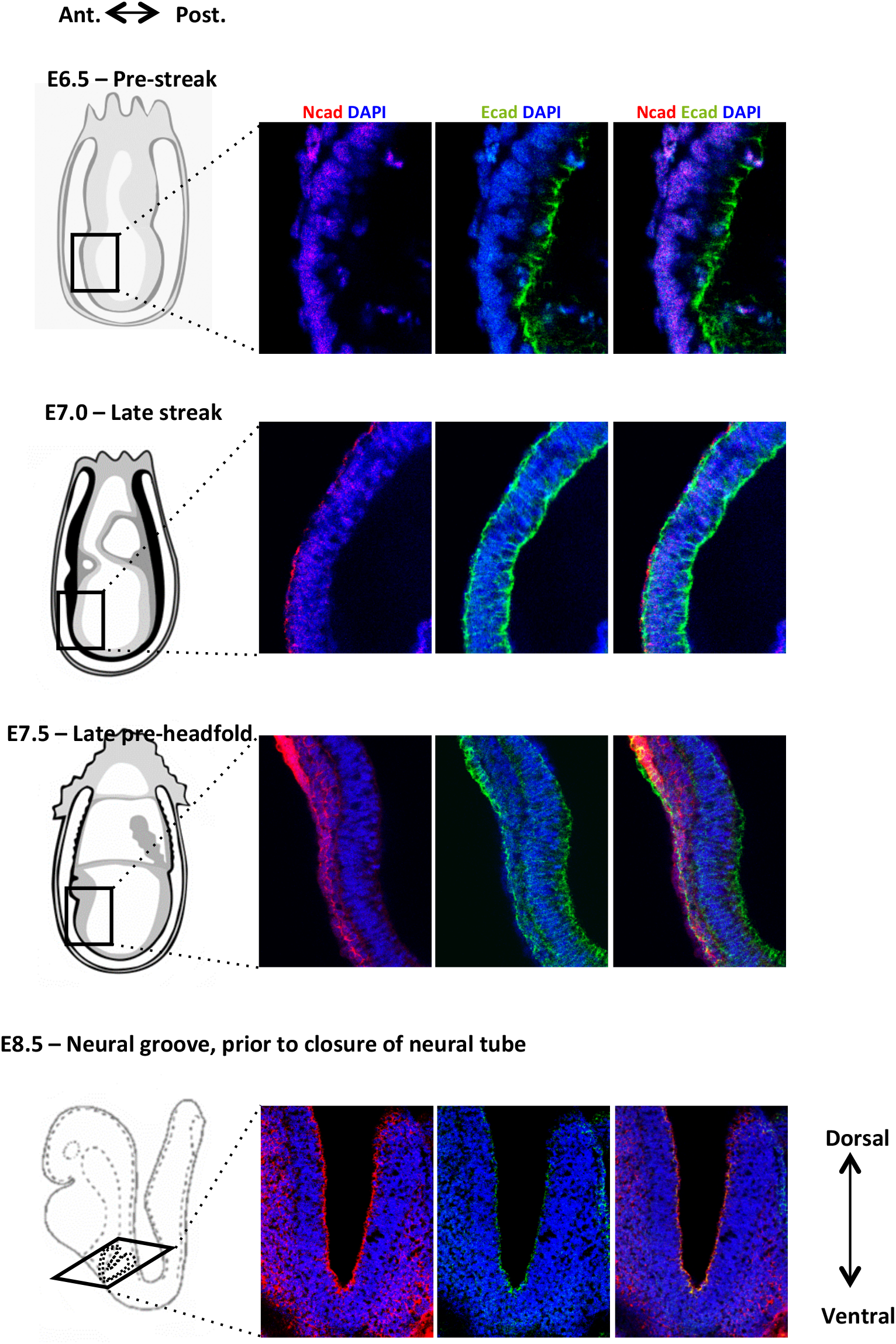
Cadherin switching in vivo. E-cadherin (green) and N-cadherin (red) immunostaining at various stages of mouse development as indicated. Embryo drawings are adapted from the EMAP eMouse Atlas Project (http://www.emouseatlas.org).

## References

1. Gilmour D, Rembold M, Leptin M. From morphogen to morphogenesis and back. Nature. 2017;541:311–20.

2. Wheelock MJ, Shintani Y, Maeda M, Fukumoto Y, Johnson KR. Cadherin switching. J Cell Sci. 2008;121(Pt 6):727–35.

3. Hatta K, Takeichi M. Expression of N-cadherin adhesion molecules associated with early morphogenetic events in chick development. Nature. 1986;320:447–9.

4. Malaguti M, Nistor P a, Blin G, Pegg A, Zhou X, Lowell S. Bone morphogenic protein signalling suppresses differentiation of pluripotent cells by maintaining expression of E-Cadherin. Elife. 2013 Jan;2:e01197.

5. Basilicata MF, Frank M, Solter D, Brabletz T, Stemmler MP. Inappropriate cadherin switching in the mouse epiblast compromises proper signaling between the epiblast and the extraembryonic ectoderm during gastrulation. Sci Rep. 2016;6(January):26562.

6. Ying Q-L, Wray J, Nichols J, Batlle-Morera L, Doble B, Woodgett J, et al. The ground state of embryonic stem cell self-renewal. Nature [Internet]. 2008 May 22;453(7194):519–23.

7. Boroviak T, Loos R, Lombard P, Okahara J, Behr R, Sasaki E, et al. Lineage-Specific Profiling Delineates the Emergence and Progression of Naive Pluripotency in Mammalian Embryogenesis. Dev Cell. 2015;35(3):366–82.

8. Tesar PJ, Chenoweth JG, Brook F a, Davies TJ, Evans EP, Mack DL, et al. New cell lines from mouse epiblast share defining features with human embryonic stem cells. Nature. 2007;448(July):196–9.

9. Brons IGM, Smithers LE, Trotter MWB, Rugg-Gunn P, Sun B, Chuva de Sousa Lopes SM, et al. Derivation of pluripotent epiblast stem cells from mammalian embryos. Nature. 2007;448(July):191–5.

10. Nichols J, Smith A. Naive and primed pluripotent states. Cell Stem Cell. Elsevier Inc.; 2009 Jun 5;4(6):487–92.

11. Chou YF, Chen HH, Eijpe M, Yabuuchi A, Chenoweth JG, Tesar P, et al. The Growth Factor Environment Defines Distinct Pluripotent Ground States in Novel Blastocyst-Derived Stem Cells. Cell. 2008;135(3):449–61.

12. Soncin F, Mohamet L, Eckardt D, Ritson S, Eastham AM, Bobola N, et al. Abrogation of E-cadherin-mediated cell-cell contact in mouse embryonic stem cells results in reversible LIF-independent self-renewal. Stem Cells. 2009 Sep;27(9):2069–80.

13. Redmer T, Diecke S, Grigoryan T, Quiroga-Negreira A, Birchmeier W, Besser D. E-cadherin is crucial for embryonic stem cell pluripotency and can replace OCT4 during somatic cell reprogramming. EMBO Rep. 2011;12(7):720–6.

14. del Valle I, Rudloff S, Carles A, Li Y, Liszewska E, Vogt R, et al. E-cadherin is required for the proper activation of the Lifr/Gp130 signaling pathway in mouse embryonic stem cells. Development. 2013 Apr;140(8):1684–92.

15. Faunes F, Hayward P, Descalzo SM, Chatterjee SS, Balayo T, Trott J, et al. A membrane-associated β-catenin/Oct4 complex correlates with ground-state pluripotency in mouse embryonic stem cells. Development. 2013;140(6):1171–83.

16. Livigni A, Peradziryi H, Sharov AA, Chia G, Hammachi F, Migueles RP, Sukparangsi W, Pernagallo S, Bradley M, Nichols J, Ko MSH, Brickman JM. A conserved Oct4/POUV-dependent network links adhesion and migration to progenitor maintenance. Curr Biol. 2013 Nov 18;23(22):2233–2244

17. Larue L, Ohsugi M, Hirchenhain J, Kemler R. E-cadherin null mutant embryos fail to form a trophectoderm epithelium. Proc Natl Acad Sci U S A. 1994;91(August):8263–7.

18. Larue L, Antos C, Butz S, Huber O, Delmas V, Dominis M, et al. A role for cadherins in tissue formation. Development. 1996;122(10):3185–94.

19. Bedzhov I, Liszewska E, Kanzler B, Stemmler MP. Igf1r signaling is indispensable for preimplantation development and is activated via a novel function of E-cadherin. PLoS Genet. 2012 Jan;8(3).

20. Zhang J, Woodhead GJ, Swaminathan SK, Noles SR, McQuinn ER, Pisarek AJ, et al. Cortical neural precursors inhibit their own differentiation via N-cadherin maintenance of beta-catenin signaling. Dev Cell. 2010 Mar 16;18(3):472–9.

21. Camus A, Perea-Gomez A, Moreau A, Collignon J. Absence of Nodal signaling promotes precocious neural differentiation in the mouse embryo. Dev Biol. 2006;295(2):743–55.

22. Di-Gregorio A, Sancho M, Stuckey DW, Crompton L a, Godwin J, Mishina Y, et al. BMP signalling inhibits premature neural differentiation in the mouse embryo. Development. 2007;134:3359–69.

23. Greber B, Wu G, Bernemann C, Joo JY, Han DW, Ko K, et al. Conserved and Divergent Roles of FGF Signaling in Mouse Epiblast Stem Cells and Human Embryonic Stem Cells. Cell Stem Cell. 2010;6:215–26.

24. Stavridis MP, Collins BJ, Storey KG. Retinoic acid orchestrates fibroblast growth factor signalling to drive embryonic stem cell differentiation. Development. 2010;137(6):881–90.

25. Sterneckert J, Stehling M, Bernemann C, Araúzo-Bravo MJ, Greber B, Gentile L, et al. Neural induction intermediates exhibit distinct roles of Fgf signaling. Stem Cells. 2010;28:1772–81.

26. Jaeger I, Arber C, Risner-Janiczek JR, Kuechler J, Pritzsche D, Chen I, et al. Temporally controlled modulation of FGF/ERK signaling directs midbrain dopaminergic neural progenitor fate in mouse and human pluripotent stem cells. Development.

27. Aubert J, Dunstan H, Chambers I, Smith A. Functional gene screening in embryonic stem cells implicates Wnt antagonism in neural differentiation. Nat Biotechnol. 2002;20(12):1240–5.

28. Haegele L, Ingold B, Naumann H, Tabatabai G, Ledermann B, Brandner S. Wnt signalling inhibits neural differentiation of embryonic stem cells by controlling bone morphogenetic protein expression. Mol Cell Neurosci. 2003;24(3):696–708.

29. Williams EJ, Furness J, Walsh FS, Doherty P. Activation of the FGF receptor underlies neurite outgrowth stimulated by L1, N-CAM, and N-cadherin. Neuron. 1994;13(3):583–94.

30. Williams EJ, Williams G, Howell F V., Skaper SD, Walsh FS, Doherty P. Identification of an N-cadherin Motif that Can Interact with the Fibroblast Growth Factor Receptor and Is Required for Axonal Growth. J Biol Chem. 2001;276(47):43879–86.

31. Utton M a, Eickholt B, Howell F V, Wallis J, Doherty P. Soluble N-cadherin stmulates fibroblast growth factor receptor dependent neurite outgrowth and N-cadherin and the fibroblast growth factor receptor co-cluster in cells. J Neurochem. 2001;76:1421–30.

32. Takehara T, Teramura T, Onodera Y, Frampton J, Fukuda K. Cdh2 stabilizes FGFR1 and contributes to primed-state pluripotency in mouse epiblast stem cells. Sci Rep. 2015;5(April):14722.

33. Howard S, Deroo T, Fujita Y, Itasaki N. A positive role of cadherin in wnt/B-catenin signalling during epithelial-mesenchymal transition. PLoS One. 2011;6(8).

34. Pieters T, van Roy F. Role of cell-cell adhesion complexes in embryonic stem cell biology. J Cell Sci. 2014 Jun 15;127(Pt 12):2603–13.

35. Dady A, Blavet C, Duband J. Timing and Kinetics of E- to N-Cadherin Switch During Neurulation in the Avian Embryo. Dev Dyn 2012;(June):1333–49.

36. Tsakiridis A, Huang Y, Blin G, Skylaki S, Wymeersch F, Osorno R, et al. Distinct Wnt-driven primitive streak-like populations reflect in vivo lineage precursors. Development. 2014 Mar;141(6):1209–21.

37. Ying Q-L, Stavridis M, Griffiths D, Li M, Smith A. Conversion of embryonic stem cells into neuroectodermal precursors in adherent monoculture. Nat Biotechnol. 2003;21(February):183–6.

38. Wood HB, Episkopou V. Comparative expression of the mouse Sox1, Sox2 and Sox3 genes from pre-gastrulation to early somite stages. Mech Dev. 1999;86:197–201.

39. Kan NG, Stemmler MP, Junghans D, Kanzler B, de Vries WN, Dominis M, et al. Gene replacement reveals a specific role for E-cadherin in the formation of a functional trophectoderm. Development. 2007 Jan;134(1):31–41.

40. Pieters T, Goossens S, Haenebalcke L, Andries V, Stryjewska A, De Rycke R, et al. p120 Catenin-Mediated Stabilization of E-Cadherin Is Essential for Primitive Endoderm Specification. PLoS Genet. 2016;12(8):1–28.

41. Libusova L, Stemmler MP, Hierholzer A, Schwarz H, Kemler R. N-cadherin can structurally substitute for E-cadherin during intestinal development but leads to polyp formation. Development. 2010;137(14):2297–305.

42. Blin G, Sadurska D, Migueles RP, Chen N, Watson JA, Lowell S. NesSys: a novel method for accurate nuclear segmentation in 3D. PLoS Biol. 2019 (in press)

43. Kalkan T, Smith A. Mapping the route from naive pluripotency to lineage specification. Philos Trans R Soc Lond B Biol Sci. 2014;369(1657):20130540-.

44. Fedor-Chaiken M, Hein PW, Stewart JC, Brackenbury R, Kinch MS. E-cadherin binding modulates EGF receptor activation. Cell Commun Adhes. 2003;10(2):105–18.

45. Wetering M Van De, Barker N, Harkes IC, Wetering M Van De, Barker N, Harkes IC, et al. Mutant E-cadherin Breast Cancer Cells Do Not Display Constitutive Wnt Signaling Mutant E-cadherin Breast Cancer Cells Do Not Display Constitutive Wnt Signaling. Cancer Res 2001;(53):278–84.

46. Hendriksen J, Jansen M, Brown CM, van der Velde H, van Ham M, Galjart N, et al. Plasma membrane recruitment of dephosphorylated beta-catenin upon activation of the Wnt pathway. J Cell Sci. 2008;121(11):1793–802.

47. Clevers H. Wnt/B-Catenin Signaling in Development and Disease. Cell. 2006;127(3):469–80.

48. Radice GL, Rayburn H, Matsunami H, Knudsen KA, Takeichi M, Hynes RO. Developmental defects in mouse embryos lacking N-cadherin. Dev Biol. 1997;181(1):64–78.

49. Orsulic S, Huber O, Aberle H, Arnold S, Kemler R. E-cadherin binding prevents beta-catenin nuclear localization and beta-catenin/LEF-1-mediated transactivation. J Cell Sci. 1999;112 (Pt 8:1237–45.

50. Cambray N, Wilson V. Two distinct sources for a population of maturing axial progenitors. Development. 2007;134(15):2829–40.

51. Takemoto T, Uchikawa M, Kamachi Y, Kondoh H. Convergence of Wnt and FGF signals in the genesis of posterior neural plate through activation of the Sox2 enhancer N-1. Development 2005;297–306.

52. Turner D a, Hayward PC, Baillie-Johnson P, Rué P, Broome R, Faunes F, et al. Wnt/β-catenin and FGF signalling direct the specification and maintenance of a neuromesodermal axial progenitor in ensembles of mouse embryonic stem cells. Development. 2014 133;141(22):4243–53.

53. Martyn I, Brivanlou AH, Siggia ED. A wave of WNT signalling balanced by secreted inhibitors controls primitive streak formation in micropattern colonies of human embryonic stem cells. Development. 2018; 146: dev172791 doi: 10.1242/dev.172791

54. Richardson L, Venkataraman S, Stevenson P, Yang Y, Moss J, Graham L, et al. EMAGE mouse embryo spatial gene expression database: 2014 update. Nucleic Acids Res. 2014;42(November 2013):835–44.

55. Smith AG. Culture and differentiation of embryonic stem cells. J Tissue Cult Methods. 1991;13(330):89–94.

56. Aubert J, Stavridis MP, Tweedie S, O’Reilly M, Vierlinger K, Li M, et al. Screening for mammalian neural genes via fluorescence-activated cell sorter purification of neural precursors from Sox1-gfp knock-in mice. Proc Natl Acad Sci U S A. 2003;100 Suppl(90001):11836–41.

57. Derksen PWB, Liu X, Saridin F, Gulden H Van Der, Zevenhoven J, Evers B, et al. Somatic inactivation of E-cadherin and p53 in mice leads to metastatic lobular mammary carcinoma through induction of anoikis resistance and angiogenesis. Cancer Cell. 2006;(November):437–49.

58. Iacovino M, Bosnakovski D, Fey H, Rux D, Bajwa G, Mahen E, et al. Inducible cassette exchange: a rapid and efficient system enabling conditional gene expression in embryonic stem and primary cells. Stem Cells. 2011;29(10):1580–8.

59. Iacovino M, Roth ME, Kyba M. Rapid genetic modification of mouse embryonic stem cells by inducible cassette exchange recombination. Methods Mol Biol. 2014;1101:339–51.

60. Iacovino M, Hernandez C, Xu Z, Bajwa G, Prather M, Kyba M. A conserved role for Hox paralog group 4 in regulation of hematopoietic progenitors. Stem Cells Dev. 2009;18(5):783–92.

61. Pollard SM, Benchoua A, Lowell S. Neural Stem Cells, Neurons, and Glia. Methods Enzymol. 2006;418(06):151–69.

62. Macleod KG, Serrels B, Carragher NO. Reverse Phase Protein Arrays and Drug Discovery. Methods Mol Biol. 2017;1647:153–69.

63. Huang DW, Sherman BT, Lempicki RA. Bioinformatics enrichment tools: paths toward the comprehensive functional analysis of large gene lists. Nucleic Acids Res. 2009;37(1):1–13.

64. Huang DW, Sherman BT, Lempicki RA. Systematic and integrative analysis of large gene lists using DAVID bioinformatics resources. Nat Protoc. 2008;(2).

